# STING agonists in combination with epigenetic drugs potentiate ZNFX1-driven inflammatory necroptosis in *TP53*-mutated AML

**DOI:** 10.64898/2026.07.27.741029

**Authors:** Kaushlendra Tripathi, Lora Stojanovic, Zahra Gohari, Mostafa Abdelrheem, Gabriella Santos, Aviva Tyler, Brandon Cooper, Rena G. Lapidus, Darren Perkins, Alonso Heredia, Kenneth P. Nephew, Stephen B. Baylin, Michael J. Topper, Maria R. Baer, Feyruz V. Rassool

**Affiliations:** Department of Radiation Oncology, University of Maryland School of Medicine, Baltimore, MD 21201, USA; University of Maryland Marlene and Stewart Greenebaum Comprehensive Cancer Center, Baltimore, MD 21201, USA; Department of Medicine, University of Maryland School of Medicine, Baltimore, MD 21201, USA; Department of Microbiology and Immunology, University of Maryland School of Medicine, Baltimore, MD 21201, USA; Institute of Human Virology, University of Maryland School of Medicine, Baltimore, MD 21201, USA; Medical Sciences Program, Indiana University School of Medicine, Bloomington, IN 47405, USA; Department of Oncology, The Sidney Kimmel Comprehensive Cancer Center at Johns Hopkins, Baltimore, MD 21231, USA; Van Andel Research Institute, Grand Rapids, MI 49503, USA; Veterans Affairs Medical Center, Baltimore, MD 21201, USA; Department of Microbiology and Immunology, VCOM Carolina Campus, Spartanburg, SC 29303, USA

## Abstract

*TP53*-mutated acute myeloid leukemia (AML) has dismal outcomes with current treatments and represents a critical unmet need. *TP53*-mutated AML is proposed to be susceptible to immunotherapeutic approaches but, to date, there is no established immunotherapy for this sub-group. Expression of stimulator of interferon genes (STING), a key innate immune driver that activates interferon (IFN) signaling, is decreased by epigenetic silencing or mutation in many cancers, including those with *TP53* mutations. Here, we report that response to the next-generation synthetic STING agonist C92 is potentiated in AML cell lines and primary cells with *TP53-*mutated versus wild-type (WT) cells, representing a previously undescribed vulnerability of these leukemia cells to STING small molecule therapies. Moreover, combining treatment with the DNA methyltransferase inhibitor (DNMTi) decitabine (DAC), significantly increases STING activation, with marked transcriptome-wide increase in repetitive elements (REs) and upregulation of a critical set of interferon-related genes. Cell death in *TP53* KO versus WT AML is specifically dependent on innate immune zinc finger NFX1-type containing 1 (ZNFX1) and Z-DNA-binding protein 1 (ZBP1) driving increased cleavage and activation of Receptor-Interacting-Serine/Threonine-Protein Kinase 3 (RIPK3) and mixed lineage kinase domain-like protein (MLKL), suggesting mechanisms of necroptosis. Finally, C92 and DAC combination significantly reduces leukemia burden in humanized AML mouse models, accompanied by increased immune responses, including cytokines and cytotoxic T lymphocytes in the leukemia microenvironment. These results support development of clinical trial strategies combining STING agonists with DNMTis for patients with *TP53*-mutated AML.

**Summary:**

- TP53-mutated AML potentiates effects of novel next-generation STING agonist C92, with unique allosteric and non-cyclic dinucleotide (non-CD) mechanism of action, inducing increased STING activation and cytokine release
- STING agonists and DNMTis, synergistically increase STING activation with marked transcriptome-wide increase in repetitive elements (REs) and upregulation of a critical set of interferon-related genes in *TP53*-mutated AML
- STING agonists induce necroptosis via a STING-ZNFX1-ZBP1-necroptosis axis in *TP53*-mutated AML.
- This drug combination reduces leukemia burden, activates immune responses in AML models and supports translation for high-risk AML patients.

**Statement of Translational Relevance:** This pre-clinical study identifies a novel therapeutic vulnerability in (*TP53*)-mutated acute myeloid leukemia (AML), a poor prognosis subtype with a critical unmet need. Novel next-generation STING agonist C92, with unique allosteric and non-cyclic dinucleotide (non-CD) mechanism of action, induces increased STING activation and cytokine release, compared with WT TP53 in AML cell lines and primary cells, and has superior STING activity with respect to several STING agonists currently in clinical studies. Combining C92 treatment with the DNA methyltransferase inhibitor (DNMTi) decitabine (DAC) synergistically increases STING activation, with marked transcriptome-wide increase in repetitive elements (REs) and upregulation of a critical set of interferon-related genes, driving ZNFX1-driven inflammatory necroptotic cell death. Utilizing humanized mouse models, C92 in combination with DAC significantly reduces leukemia burden and enhances cytotoxic T-cell responses in the tumor microenvironment, supporting clinical translation for high-risk AML patients.

## Introduction

Acute myeloid leukemia (AML) is a genetically heterogeneous disease(1). Mutations in *TP53,* which encodes the DNA damage response transcription factor p53, are present in approximately 10% of *de novo* and 25% of therapy-related AML at diagnosis, and are associated with poor-risk cytogenetic changes and poor outcomes following both chemotherapy and allogeneic hematopoietic stem cell transplantation (2,3). Approximately 80% of *TP53* mutations in AML are missense, occurring within the DNA-binding domain and resulting in loss of DNA-binding capability of the encoded protein(4).

DNA methyltransferase inhibitors (DNMTis) are FDA-approved for treatment of myelodysplastic syndrome (MDS) and AML. *TP53*-mutated AML cells are sensitive to DNMTis in preclinical settings(5,6) and, clinically, *TP53*-mutated AML has higher response rates to DNMTis than to chemotherapy(7). However, responses of *TP53*-mutated AML to DNMTis are not durable(8). Combination treatment with the BCL2 inhibitor venetoclax (Ven) and DNMTis has improved treatment outcomes in AML patients not eligible for chemotherapy(9), but has produced little improvement in *TP53*-mutated AML. Recent clinical data suggest that *TP53*-mutated AML may be susceptible to immunotherapeutic approaches (10,11), but none of these has an established role in AML to date. Therefore, developing immunotherapy strategies in patients with *TP53*-mutated AML remains an unmet need.

We reported that DNMTis, in combination the poly ADP-ribose polymerase inhibitor (PARPi) talazoparib increased activation of stimulator of interferon genes (STING)-dependent IFN and inflammasome signaling in AML cells with, compared to those without *TP53* mutations(12). STING plays a key role in the anticancer innate immune response(13), generating IFN signaling that increases T cell infiltration, and leading to anti-tumor immune responses. Basal STING expression is decreased in many cancers due to DNA methylation or mutation(14). STING activation is triggered by dsDNA binding to the cytosolic receptor cyclic GMP-AMP synthase (cGAS). PARPis have also been reported to activate STING-dependent IFN signaling through increased cytosolic dsDNA(15). DNMTi treatment induces “viral mimicry” with a critical role in the re-expression of hypermethylated endogenous retroviral (ERV) elements and cytosolic dsRNA that activate IFN signaling(16,17). DNMTis can also transcriptionally increase STING and thereby increase the quantity of STING available for activation(18). In this context, Diepstraten *et al*. reported that the therapeutic barrier imposed by defects in *TP53* can be overcome by direct activation of the cGAS/STING pathway, which promotes apoptosis of blood cancer cells through *TP53*-independent BH3-only protein upregulation. Combining clinically-relevant STING [agonists](https://www.sciencedirect.com/) with BH3-mimetic drugs efficiently kills *TRP53*/*TP53*-mutant mouse B lymphoma, human NK/T lymphoma and AML cells (19). Taken together, the data suggest that STING agonists in combination with mechanistically-relevant targeted therapeutics, including BH3-mimetic drugs, could be a promising therapy strategy to treat patients with *TP53*-mutant blood cancers(20).

First-generation STING agonists have been in clinical trials in solid tumors, but responses have been suboptimal, likely due to targeting of mainly mouse rather than human STING as well as poor delivery capability, requiring intra-tumoral administration (21,22). Importantly, in this regard, Curadev Pharma has developed a potent next-generation STING agonist, C92, that is specific for human STING and can be administered intravenously (23). This drug binds all variants of human STING and causes allosteric changes that stabilize the activated dimer (24) and promises to bridge the translational gap in STING agonist immunotherapy, showing that it delivers strong IFN-I responses with minimal non-IFN pharmacology by blocking STING’s proton channel activity (23). Phenotypically, this results in antitumor immune responses unimpeded by immune tachyphylaxis and heightened levels of systemic tolerability for the treatment of patients with advanced/metastatic cancer. Studies presented also demonstrate that STING activation by C92 significantly retards the growth of high chromosomal instability tumors in mice (23).

We report in this pre-clinical study, a novel therapeutic vulnerability in (*TP53*)-mutated acute myeloid leukemia (AML), a poor prognosis subtype with a critical unmet need. C92 induces increased STING activation and cytokine release, compared with WT *TP53* in AML cell lines and primary cells, and has superior STING activity with respect to several STING agonists currently in clinical studies. Combining C92 and DNMTis produces synergistic cytotoxicity that is mediated by the master innate immune regulators, ZNFX1 and ZBP1, leading to necroptotic cell death. In studies in immune-competent AML models, this is accompanied by increases in immune responses, including cytokines and cytotoxic T lymphocytes in the tumor microenvironment. These studies provide new and translationally relevant insights into the importance of loss of *TP53* in STING activation and transcriptional activation of repeat elements and interferon-driven cell death and antileukemia immune responses in AML with *TP53* mutations. The data also suggests potential treatment strategies for other *TP53*-mutated cancers, in addition to *TP53*-mutated AML.

## Materials and methods

### Cell lines

AML cell lines MOLM-14 (RRID:CVCL_7916), KG-1a (RRID:CVCL_1824), U937 (RRID:CVCL_0007) and Kasumi-1 (RRID:CVCL_0589) were cultured in RPMI 1640 (Corning) supplemented with 10% FBS (Sigma) and 1% penicillin-streptomycin (Sigma) at 37°C in 5% CO_2_. MOLM-14. To ensure cell line integrity, cell lines were thawed at frequent intervals and were tested regularly for mycoplasma.

### CRISPR CAS9 KO

CRISPR cell lines with knockout (KO) of the *TP53* and/or *ZNFX1* genes were generated in the Translational Laboratory Shared Resources CRISPR Core (TLSR-CRISPR) using the CRISPR-Cas9 mechanism. Synthetic single-guide RNAs (sgRNAs, Synthego) were generated targeting *TP53* exons 4 and 6 and *ZNFX1* exons 8 and 11. CRISPR-Cas9 KOs were produced by nucleofection on the Lonza Amaxa™ 4D-Nucleofector platform and confirmed by subjecting cells to PCR and Sanger sequencing. Genomic editing was confirmed by INDEL analysis using the Synthego ICE analysis platform. Clonal KO populations were then generated by single-cell plating and screening of clonal sequences using ICE analysis.

### sgRNA Sequences

TP53 Exon 4: CCATTGTTCAATATCGTCCG

TP53 Exon 6: CAGACCTCAGGCGGCTCATA

ZNFX1 Exon 8: GCCATGAGGCTAGACCATTG

ZNFX1 Exon 11: CATTTTGATTGGGGACCACC

### Primary AML cells

Blood samples were obtained from AML patients with peripheral blasts following informed consent on a protocol approved by the University of Maryland School of Medicine (UMSOM) Institutional Review Board. Mononuclear cells (MNCs) were isolated by density centrifugation over Ficoll-Paque (Sigma-Aldrich) and were cultured in IMDM with 20% FBS, without cytokine supplementation, at 37°C in 5% CO_2_.

### *in vitro* treatments

5-aza-2-deoxycytidine (decitabine/DAC) (MedChem Express) was prepared at 5mM in DMSO and aliquoted and stored at -80°C for single use. STING agonist C92 (Curadev Pharma, city, country) was prepared at 5mM in DMSO. Caspase inhibitor Z-VAD (Sigma-Aldrich) was prepared at 10mM in DMSO and used in concentration of 20µM. Janus kinase (JAK) inhibitor ruxolitinib was prepared at 10mM in DMSO and used at concentration of 10µM. TBK1/IKKe inhibitor amlexanox, was prepared at 10mM in DMSO and used at 10µM concentration. The STING agonists ADU-S100, MSA-2, diABZI were prepared at 5mM in DMSO and used at 1µM concentration. *in vitro* treatments were performed as indicated in the text, with mock treatments with DMSO at identical final concentrations.

### MTS and Synergy assay

MOLM-14, MOLM-14 *TP53* KO, U937 and Kasumi-1 cells were seeded into 96-well plates at 1 × 10⁴ cells per well in 100 µL total volume. Cells were treated daily for 3 consecutive days with C92 and DAC at varying concentrations. For MOLM-14 cells, C92 was tested in combination 10, 50, and 125 nM DAC. For U937 cells, C92 was tested at 10, 5, and 2.5 µM in combination with 1000, 100, and 10 nM DAC. After 72 hours, 20 µL CellTiter 96® AQueous One Solution Cell Proliferation Assay reagent (Promega, G3580) was added to each well. Absorbance was measured after 1 hour using a VersaMax microplate reader controlled by SoftMax Pro 6.5.1 software. Drug synergy was analyzed using the SynergyFinder Plus (RRID:SCR_019318) web platform(25). Synergistic interactions were evaluated based on the Highest Single Agent (HSA) model. In addition, combination index (CI) values were calculated using the Chou-Talalay method (26), and CI graphs were generated using GraphPad Prism 11.

### Quantitative real-time PCR (qRT-PCR)

Total RNA was isolated for qRT-PCR analysis to measure mRNA abundance of the genes indicated in **Table 2**, normalized to beta actin. Changes in RNA expression were measured by the ΔΔCT method. Data are presented as fold change after drug, versus mock, treatment.

### Immunoblotting

Cells were lysed in RIPA buffer (Sigma) supplemented with 1% phosphatase and 1% protease inhibitors (Thermo Scientific). After whole cell lysate collection and the bicinchoninic acid assay (BCA, Thermo Scientific) protein quantification, 20 µg of whole cell protein was loaded onto 4-20% SDS-PAGE gel (Bio-Rad) and transferred to polyvinylidene difluoride membranes (GE Life Science). Membranes were blocked with 5% nonfat milk before overnight incubation at 4°C with antibodies including pTBK1 (1:1000, Cell Signaling Technology, Cat# 5483, RRID:AB_10693472), STING (1:1000, Cell Signaling Technology, Cat# 13647, RRID:AB_2732796 and Cat# 50494, RRID:AB_2799375), p-STING (1:1000, Cell Signaling Technology, Cat# 19781, RRID:AB_2737062 and Cat# 85735, RRID:AB_2801279), IRF3 (1:1000, Cell Signaling Technology, Cat# 4302, RRID:AB_1904036), p-IRF3 (1:1000, Cell Signaling Technology, Cat#4947, RRID:AB_823547), ZNFX1 (1:1000 Abcam, ab179452, RRID:AB_3073968), ZBP1 (1:1000 Abcam, ab232977, RRID:AB_2911299), p53 (1:1000 Cell Signaling Technology, Cat# 18032, RRID:AB_2798793) and beta-tubulin (1:1000, Cell Signaling Technology, Cat# 86298, RRID:AB_2715541 and Cat# 2128, RRID: AB_823664). Blots were then washed three times with Tris Buffered Saline with Tween 20 (TBS-T) and then incubated with horseradish peroxidase (HRP)-conjugated secondary antibodies [HRP anti-rabbit (Cat# 111-035-003, RRID:AB_2313567) or anti-mouse (Cat# 115-035-003i, RRID:AB_10015289), Jackson Immuno-chemicals]. Blots were developed following incubation with the ECL Plus Western Blotting Detection Reagent (Amersham Bioscience) or using a Hi/Lo Digital-ECL Western Blot Detection Kit (Kwik Quant) and Kwik Quant Imager in the 1:10 ratios. Band densities were quantified using ImageJ (RRID:SCR_003070) software.

### Measurement of cytokine release by ELISA

Cytokine release was measured in both *TP53* wild-type (WT) and *TP53* KO AML cells treated in triplicate with DAC and C92 at different concentrations, and ELISA was performed. Absorbance was taken at 450 nm using a VersaMax ELISA Microplate Reader (Molecular Devices). IFN-γ, TNF-α, CXCL10, IFI27 were measured using human IFN-γ ELISA (Invitrogen), human TNF-α ELISA (Invitrogen), human CXCL10/IP-10 ELISA (Bio-Techne), and Human IFI27 ELISA (R&D System) kits.

### Apoptosis analysis

Approximately 2.5×10^5^ cells were harvested and stained with Annexin V-FITC Apoptosis Staining Kit (ab14085). Cells were incubated at room temperature in the dark for 5 minutes with Annexin V and propidium iodine (PI 50µg/ml), after which Annexin V-FITC binding was assessed by flow cytometry (Ex = 488 nm; Em = 530 nm) using FITC signal detector and PI staining by the phycoerythrin emission signal detector. Results were analyzed using Microsoft Excel (RRID:SCR_016137). and GraphPad Prism (RRID:SCR_002798).

### Humanized mouse studies

All mouse studies were approved by the University of Maryland School of Medicine (UMSOM) Institutional Animal Care and Use Committee (IACUC), and all animals were treated in accordance with the NIH guidelines for Laboratory Animals and in compliance with federal regulations, policies, and guidelines, including the Animal Welfare Act and the Public Health Service Policy. Mice were housed in microinsulator cages in Specific Pathogen Free (SPF) rooms. NSG-SGM3 (RRID:IMSR_JAX:013062) mice (NOD.Cg-*Prkdc^scid^ Il2rg^tm1Wjl^* Tg (CMV-IL3, CSF2, KITLG)1Eav/MloySzJ) were from Jackson Laboratories (Strain #013062)(27). Human CD34+ cells, isolated from cord blood, were purchased from Lonza (Walkersville, MD) or the New York Blood Bank (New York, New York). For generation of humanized CD34 mice, newborn mice from strain NSG-SGM3 underwent 62 cGy total body radiation in an RS-2000 x-ray radiator, followed by intrahepatic injection of 10^5^ CD34+ cells under anesthesia as described (28). On week 12 after transplantation, mice were checked for human cell reconstitution by double staining with FITC-conjugated anti-human CD45 antibody (BD Pharmingen, Cat# 560976, RRID:AB_395874) and APC-conjugated anti-mouse CD45 antibody (BD Pharmingen, Cat# 553079, RRID:AB_394609). Humanized mice are designated as CD34-NSG-SGM3. Assignment to experimental groups was performed randomly after successful humanization.

### Immunohistochemistry

Spleens excised from mice after euthanasia were fixed overnight in 10% formalin, embedded in paraffin, and sectioned. The sections underwent immunohistochemical staining using routine methods. Briefly, sections (5 μm) were deparaffinized, endogenous peroxidase was inactivated in 3% peroxide for 10 min, and antigen retrieval in 0.1 M sodium citrate was performed in a pressure cooker before the sections were blocked with 5% BSA and incubated overnight at 4 °C with polyclonal antibodies against CXCL10/IP-10 (Cat# 10937-1-AP, RRID:AB_2088002) and CD8 (Mouse anti-Human CD8 Antibody, Cat# 188-10258-IHC, RayBiotech, RRID:AB_2857849). CXCL10 and CD8 primary antibodies were detected using SignalStain® Boost Detection Reagent (Rabbit: Cat# 8114, RRID:AB_10544930, Mouse: Cat# 8125, RRID:AB_10547893) and developed with SignalStain® DAB Substrate Kit, then dehydrated with alcohol solutions and mounted. Slides were imaged with a Motic EasyScan scanner and analyzed with QuPath (RRID:SCR_018257) software.

### Transcriptome profiling with RNA Seq

#### Library construction and sequencing

RNA was isolated from cell pellets using the RNeasy kit (Qiagen) according to manufacturer’s instructionsQuality and quantity of RNA were measured using the Bioanalyzer RNA High Sensitivity kit on a Model 2100 BioAnalyzer (Agilent). All RNA samples had a RIN score of 9 or greater. For RNA sequencing experiments, 500 ng of total RNA was used for library construction with the TruSeq Stranded Total RNA Library kit (Illumina) according to manufacturer’s instructions. Quality and quantity of the resulting cDNA were measured using the Bioanalyzer DNA High Sensitivity kit (Agilent). Bulk RNA libraries were sequenced on an Illumina Novaseq X instrument using 150bp paired-end dual indexed reads and 1% of PhiX control. Depth of coverage was targeted at a total of 50 million reads (25 million pairs) per library.

### Bioinformatics analysis

RNA-seq analyses were conducted as follows: Raw FASTQ files were first assessed for quality metrics using FastQC (RRID:SCR_014583) followed by processing using Trimmomatic (RRID:SCR_011848) (29) to remove adapters and low-quality reads. Processed FASTQ files were then loaded into the Salmon (30) and processed, GENCODE transcript fasta was used as transcript reference. Salmon-processed files were used as input for tximport (31) followed by DESeq2 (RRID:SCR_000154) (32) for differential expression analysis and derivation of VST normalized counts. Volcano plots of differential expression data were generated using EnhancedVolcano. Pathway analyses were conducted on VST normalized counts using fgsea and specifically utilizing the GESECA workflow. Heatmaps were generated using heatmap and utilized VST normalized counts as input. For repetitive element analysis, TEtranscripts was run as previously defined (33) with Hammell lab curated GTF (GRCh38_GENCODE_rmsk_TE.gtf) used for annotation. Raw and processed data derived from these RNA-seq studies are publicly deposited in GEO (GSE327164).

### Statistical Analysis

All data are presented as mean ± SEM with statistical significance derived from two-tailed unpaired Student’s t-test using Microsoft Excel (RRID:SCR_016137). or ANOVA in GraphPad Prism, version 10.0.0 (GraphPad Software, RRID:SCR_002798).

### Data Availability

The RNA-seq data from AML cells treatments generated in this study are publicly available in GEO at GSE327164. All other raw data generated in this study are available upon request from the corresponding author.

## Results

We previously reported that DAC in combination with the PARPi talazoparib (TAL) activates STING signaling in AML cells in a *TP53*-independent manner(12). Therefore, we hypothesized that treatment with STING agonists could potently activate downstream IFN and inflammasome signaling, leading to anti-leukemia immune responses, especially in *TP53*-mutated AML. To test our hypothesis, we used the potent next-generation STING agonist C92 developed by Curadev Pharma. C92 and its analogues have unique chemical properties, binding outside of the canonical nucleotide binding pocket, which is targeted by other synthetic activators, and causing allosteric changes that stabilize the activated dimer (24). C92 is also specific for all variants of human STING and not mouse STING(34).

Our first important finding is that STING agonist C92 activates STING in a TP53*-*independent manner in AML cells. MOLM-14 AML cells treated with 100-250 nM of C92 for 3 days induce proinflammatory cytokines, including CXCL10, IFN β, and IRF3 (**SFig 1A-C**), as measured by ELISA assays, and activation of STING (phospho-STING) and its downstream target, TBK1, within 60 minutes by western blotting analysis (**Fig 1A**). However, CRISPR KO of *TP53* in MOLM-14 AML demonstrates significantly increased STING activation **(Fig 1A, SFig1D).** This *TP53*-independent effect of C92 on STING activity is generalized to other synthetic STING agonists currently in early clinical trial settings, including ADU-S100, MSA-2 and diABZl (21,22), as they also exhibit this potentiation in *TP53* KO MOLM-14 AML cells, compared with MOLM-14 parental cells, with cytokine activation measured by ELISA assays (**Fig 1B-E**). Notably, C92 demonstrates superior increase of interferon stimulated protein IFI27 and induction of cytokines TNFα, IFNγ, and CXCL10, compared with other STING agonists tested. C92 treatment also induces similarly high levels of cytokines (CXCL10, TNFα, IFNγ γ) in AML cell lines with naturally occurring *TP53* mutations, including Kasumi (R248 mutation), KG1a (defective donor splice site in intron 6) and U937 (Val173Trpfs*59) **(Fig 1F-H)**. C92 treatment also induces similar increases in cytokine expression in primary AML samples with *TP5*3 mutations (N=5), compared with WT TP53 (N=5) (**Fig. 1I-L**, **Table 1**), validating above results in AML cell lines.

**Fig 1.**
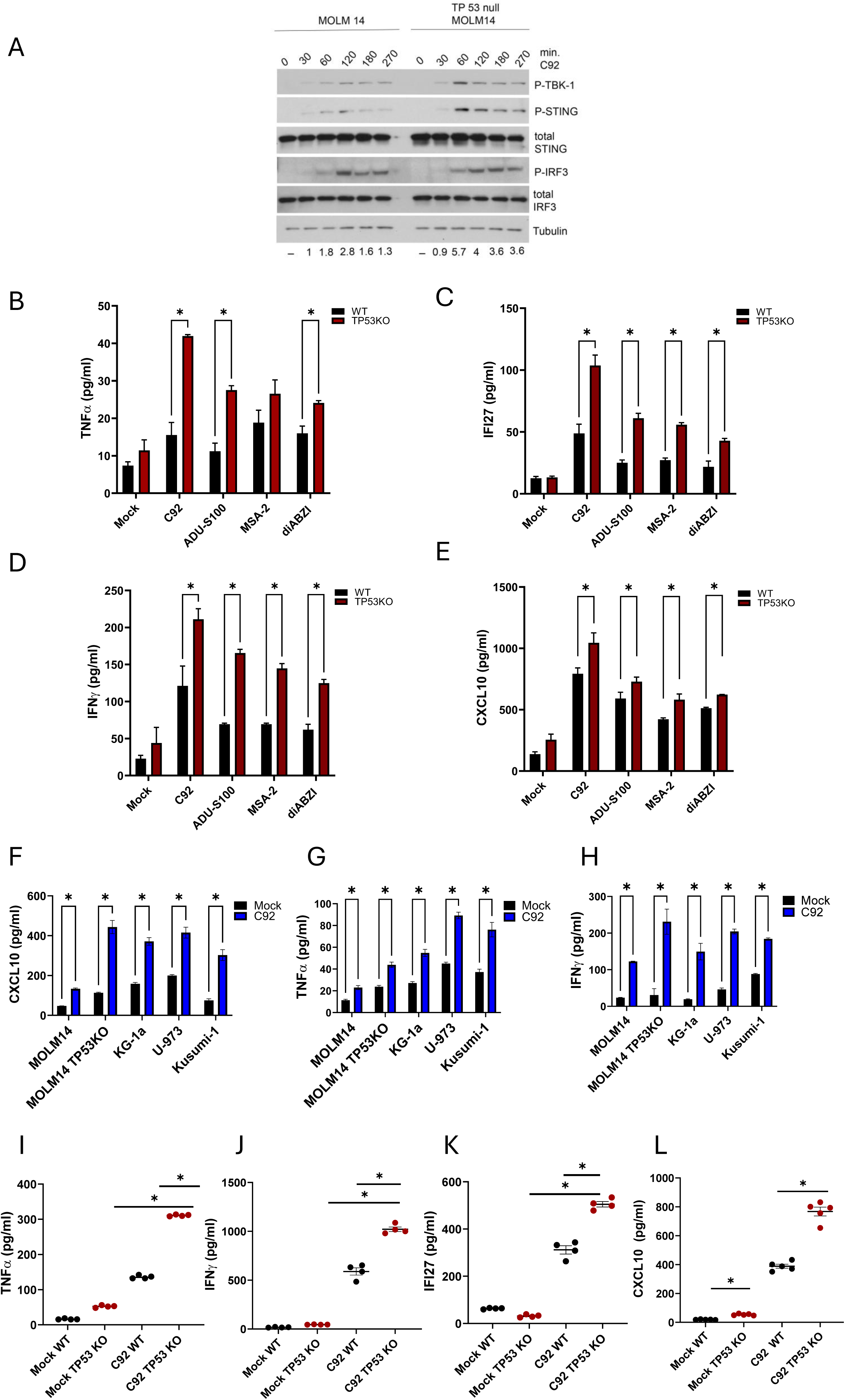
STING agonist C92 activates STING in a *TP53*-independent manner. **A.** Western blot showing STING pathway activation through different time points during C92 drug treatment. Quantitation of phospho STING with respect to loading control is shown below. **B-E.** TNF alpha, IFI27, IFN gamma, CXCL10 respectively cytokine expression measured by ELISA. **F-H**. TNF alpha, CXCL10 and IFN gamma respectively cytokine expression in different AML cell lines with p53 naturally occurring mutation. **I-L.** Cytokine expression TNF alpha, IFN gamma, IFI27 and CXCL10 respectively studied in primary (N=5) cells with TP53 mutation after the C92 treatment and measured by ELISA. All data are presented as mean +/- SEM with p-values derived from two-tailed unpaired Student’s t test or ANOVA as appropriate. * p<0.05, p<0.01, p<0.001, p<0.0001. All experiments were performed at least 3 times.

**Table 1.** Molecular and cytogenetic features of AML patients. VAF – variant allele frequency

| Patient | Age (years)<br>Sex | WBC (x10<sup>9</sup>/L) | Blood blasts (%) | TP53 Status | Karyotype | Other mutations |
| --- | --- | --- | --- | --- | --- | --- |
| TP53 mut 1 | 79F | 19.5 | 4<br>61 monos | TP53 c.824G>A, p.Cys275Tyr (NM_000546.5) VAF: 77.7% | 43<br>46,XX,add(2)(q36),t((3;6)(p21;p23),-5,add(10)(p15),-11,-13,-13,-14,add(16)(q21),<br>-17,-22,+4<br>8mar[cp11]/46,XX[9] | TET2 c.4813G>T, p.Ala1605Ser (NM_001127208.2) VAF: 47.8% |
| TP53 mut 2 | 61F | 56.4 | 70 | TP53 p.Tyr236Cys c. 707A>G 97% | 42,X,der(X;17)(q10;p10),-5,der(11)add(p15)hsr(11)(q23)add(11)(q23)hsr(11)(q23),add(12)(p12),-13,add(15)(q15),add(15)(p11.2),add(19)(p13),-22,+mar[cp14]/42,idem,r(11)add(p15)hsr(11)(q23)add(11)(q23)hsr(11)(q23),-der(11)[1] | None |
| TP53 mut 3 | 48F | 46.3 | 58 | TP53 c.423C>G, p.Cys141Trp (NM_000546.5) VAF: 46.3%<br>TP53 c.818G>A, p.Arg273His (NM_000546.5) VAF: 45.0% | 43-45,XX,+1,add(1)(q43),del(5)(q13q31),+6,-7,-16,-18,-21,der(22)ins(22:?) (q11.2;?)dup(21) (q11.2q13),+1-2mar[cp20] | NRAS c.35G>A, p.Gly12Asp (NM_002524.4) VAF: 11.1%<br>NRAS c.35G>T, p.Gly12Val (NM_002524.4) VAF: 2.8% |
| TP53 mut 4 | 67F | 37.7 | 55 | Gene Mutated: TP53<br>Mutation Site: p.A276D<br>Mutation Type: Point<br>Mutation Level: 85%<br>Exon: 8<br>Nucleotide Change: c.827C>A<br>Reference to Mutation: NONE<br>Reference to Mutation (2): COSM45268 | 45,XX,inv(3)(q21q26),-5,del(7)(q22),-17,+mar[19]/46,XX[1] | None |
| TP53 mut 5 | 62F | 10.8 | 19 | TP53p.(H179D) c.535C>G VAF 77% | 43,X,-X,del(5)(q22q35),-17,-18,-22,+mar[20] | None |
| TP53 WT 1 | 65M | 2.1 | 64 | - | 50,XY,+2,+8,der(10)t(10;12)(p12;q13),del(11)(q13q23),+19,+21[19]/46,XY[1] | KIT p.(V5301)C. 1588G>A VAF 50% |
| TP53 WT 2 | 38F | 100.2 | 80 | - | 46,XX[20] | NRASp.(G13D)c.38G>A VAF 20%<br>DNMT3A p.(R882H)c.2645G>A VAF 25%<br>NPM1 p.(W288fs)c.863_864insTCTG VAF 25%<br>PTPN11 p.(N58Y) c.172A>T VAF 5.4%<br>FLT3p.(N676K)c.2028C>A VAF 7.7%<br>IDH2 p.(R140Q)c.419G>A VAF45% |
| TP53 WT 3 | 63F | 58.2 | 34 | - | 46,XX,i(17)(q10)[19]/46,XX[1] | ETV6 p.(F368L)c.1102T>C VAF44%<br>SETBP1 p.(D868N)c.2602G>A VAF 50%<br>ASXL1 p.(W796fs*4) c.2385_2386ins VAF 48% |
| TP53 WT 4 | 61F | 17.8 | 44 | - | 45,XX,-3,-4,add(5)(q13),der(5)t(5;12)(p15;q13),del(6)(p12),add(7)(q11.2),add(11)(p14),<br>-12,der(14)t(8;14)(q13;p11.2),der(16)t(16;17)(q12-13;q21),-17,add(18)(q21),+2mar[cp13] | - |
| TP53 WT 5 | 26F | 12.3 | 49 | - | 46,XX,t(9;11)(p21;q23)[5]/47,idem,+der(9)t(9;11)[6] | NRAS p.(G12A) c.35G>C VAF 11% |

In preclinical settings, *TP53*-mutated AML cells show sensitivity to DNMTis(5,6). In this regard, we have reported that DNMTi treatment increases STING expression in multiple tumor types including AML(12,18). Therefore, we determined whether combining DAC with C92 treatment might increase basal levels of STING, to augment STING agonist C92 induction of STING (phospho-STING) as well as its downstream targets. Our data shows that, indeed, this strategy increases activation of STING and downstream target phospho-IRF3, by several fold in *TP53* KO AML cells (**Fig 2A**). Furthermore, both these increases are synergistic, inducing cytotoxicity, as assessed by models, Synergy finder plus(25) and CalcuSyn(26), for all examined AML cell lines examined (WT MOLM-14 or MOLM-14 TP53 KO,*TP53*-mutated-U937 and KG1a) regardless of TP53 status. As such, the synergy score (HSA) shows that the drug combination is more effective than DAC or C92 treatments alone giving Calcusyn combination indices of <1 (range) for most drug concentrations in MTS assays (**Fig 2B-E, SFig 2-4)**. Importantly, synergy is exponentially greater in MOLM-14 with KO of *TP53 [*12.62 (p=1.96e^-7^)], compared to MOLM-14 [mean 4.8 (p=2.69e^-01^) parental cells (**Fig 2D,E, SFig 2A,B**).

**Fig 2.**
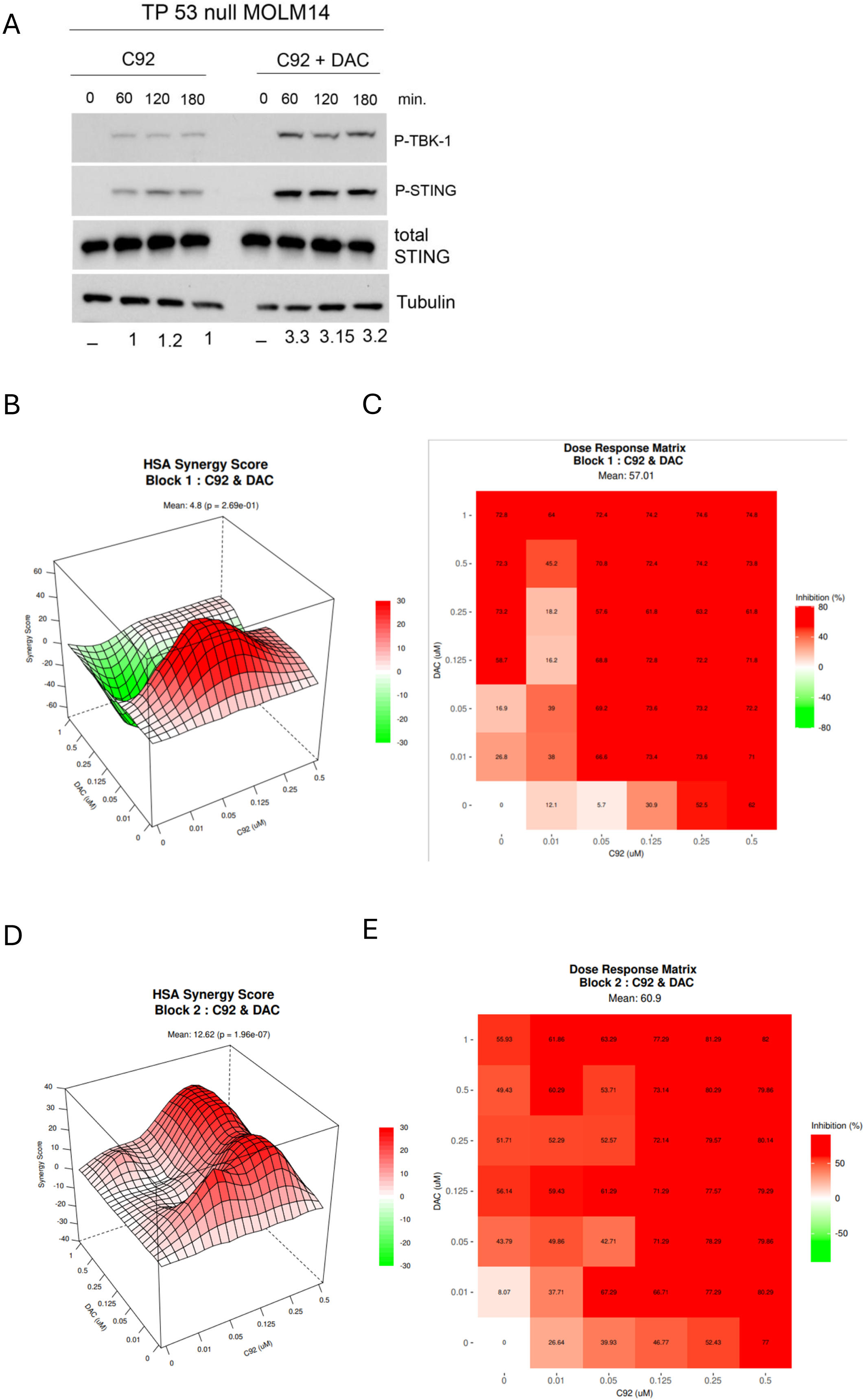
C92 and DAC treatment is synergistic in *TP53* knockout vs WT AML cells, increasing STING activation. **A.** Western blot comparing STING activation after treatment with C92 single agent and in combination with DAC in TP53 null MOLM-14 cells. Quantitation of phosphor-STING with respect to loading control is shown below. **B-E.** Cell viability assay (MTS) showing the synergistic Comparative synergy analysis between C92 and DAC in MOLM-14 WT (B,C) and *TP53* KO cells **(D,E)**. The SynergyFinderPlus tool was used to visualize drug interaction effects following exposure to a concentration matrix of decitabine (DAC) and C92 **(B,D).** Heatmap visualization and synergy scores were generated using HSA model **(C,E)**. All data are presented as mean +/- SEM with p-values derived from two-tailed unpaired Student’s t test or ANOVA as appropriate. * p<0.05, p<0.01, p<0.001, p<0.0001. All experiments were performed at least 3 times.

**Fig. 3.**
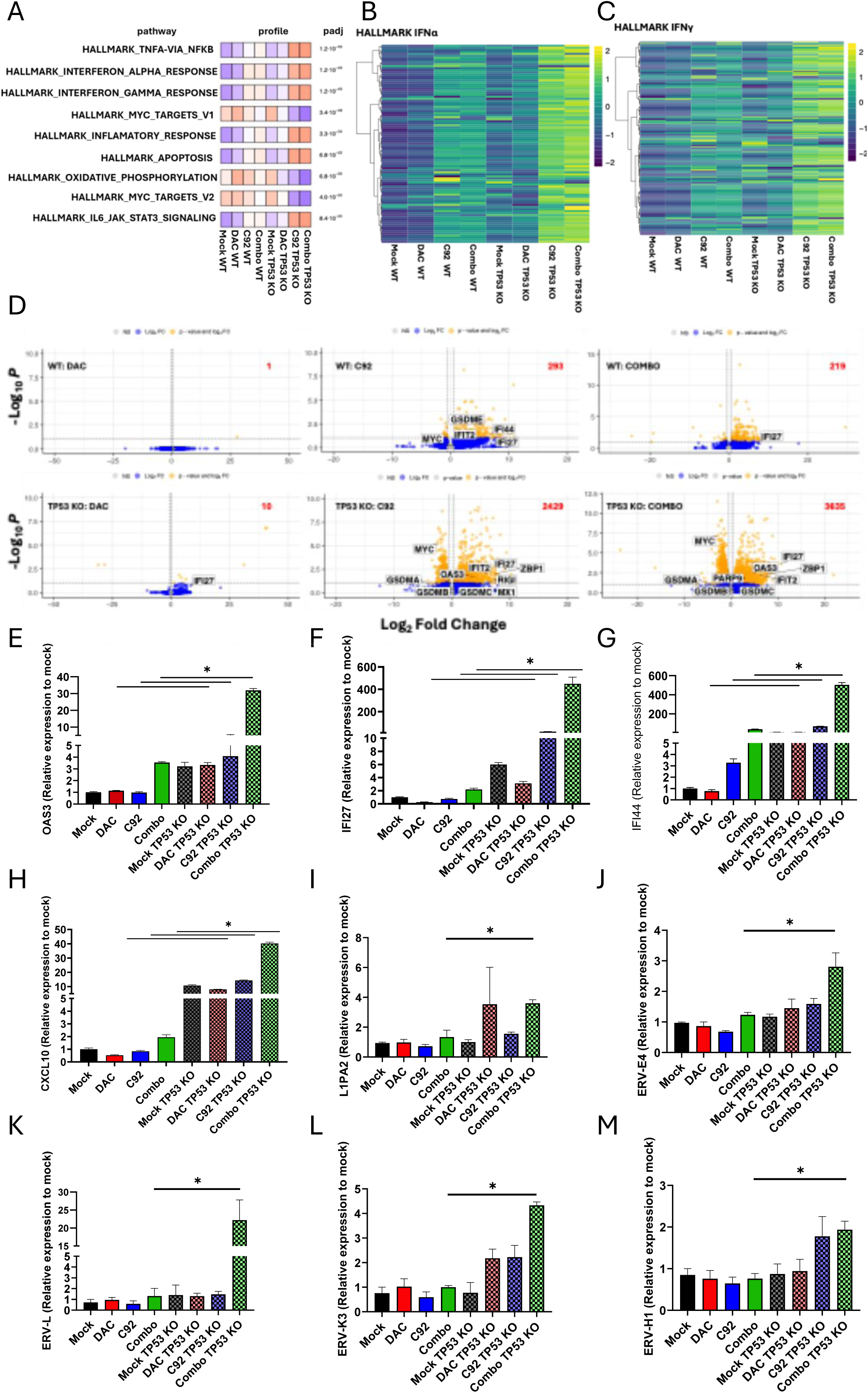
TP53 status drives transcriptional response dynamics to DAC +/- STING agonist (C92). **A**. GESCA-derived pathway analysis for msigdb HALLMARKS based on DESeq2 VST normalized RNA-seq counts, n=2 averaged RNA-seq datasets collected for MOLM-14 WT and MOLM-14 TP53 KO after 3 days treatment with C92 (250nM and DAC (25nM). Red color indicates positive enrichment, blue indicates negative enrichment, padj for pathway enrichment provided in figure. Heatmap of median VST normalized counts for (**B**) Hallmarks IFNα, (**C**) Hallmarks IFNγ. Unsupervised hierarchical clustering of genes by Ward.D2. **D.** Volcano plots for GENCODE gene symbols from n=2 averaged RNA-seq datasets for MOLM-14 WT and MOLM-14 TP53 KO after 3 days treatment with C92 (250nM and DAC (25nM). Differentially expressed genes were derived by DESeq2 pipeline and are **orange**, and the number of DEGs per condition is in **red** text in the figure. MYC and interferon associated genes are labeled within plot. **E-H.** Bar graph showing relative transcript levels of interferon gene categories **E.** CXCL10, **F.** IFI227, G. IFI44L, and **H.** OAS3 in MOLM-14 WT and MOLM-14 TP53 -/- cells following 3 days treatment with 2.5nM DAC, 100nM C92, or combination. **I-M.** Bar graph showing RE relative transcript levels **I.** L1PA2, **J.** ERV-E4, **K.** ERV-L, **L**. ERV-K3, and **M.** ERV-H1 performed by qPCR following 3 days of treatment with DAC (10nM) and C92 (100nM). All data are presented as mean +/- SEM with p-values derived from two-tailed unpaired Student’s t test or ANOVA as appropriate. * p<0.05, p<0.01, p<0.001, p<0.0001. All experiments were performed at least 3 times.

**Fig. 4.**
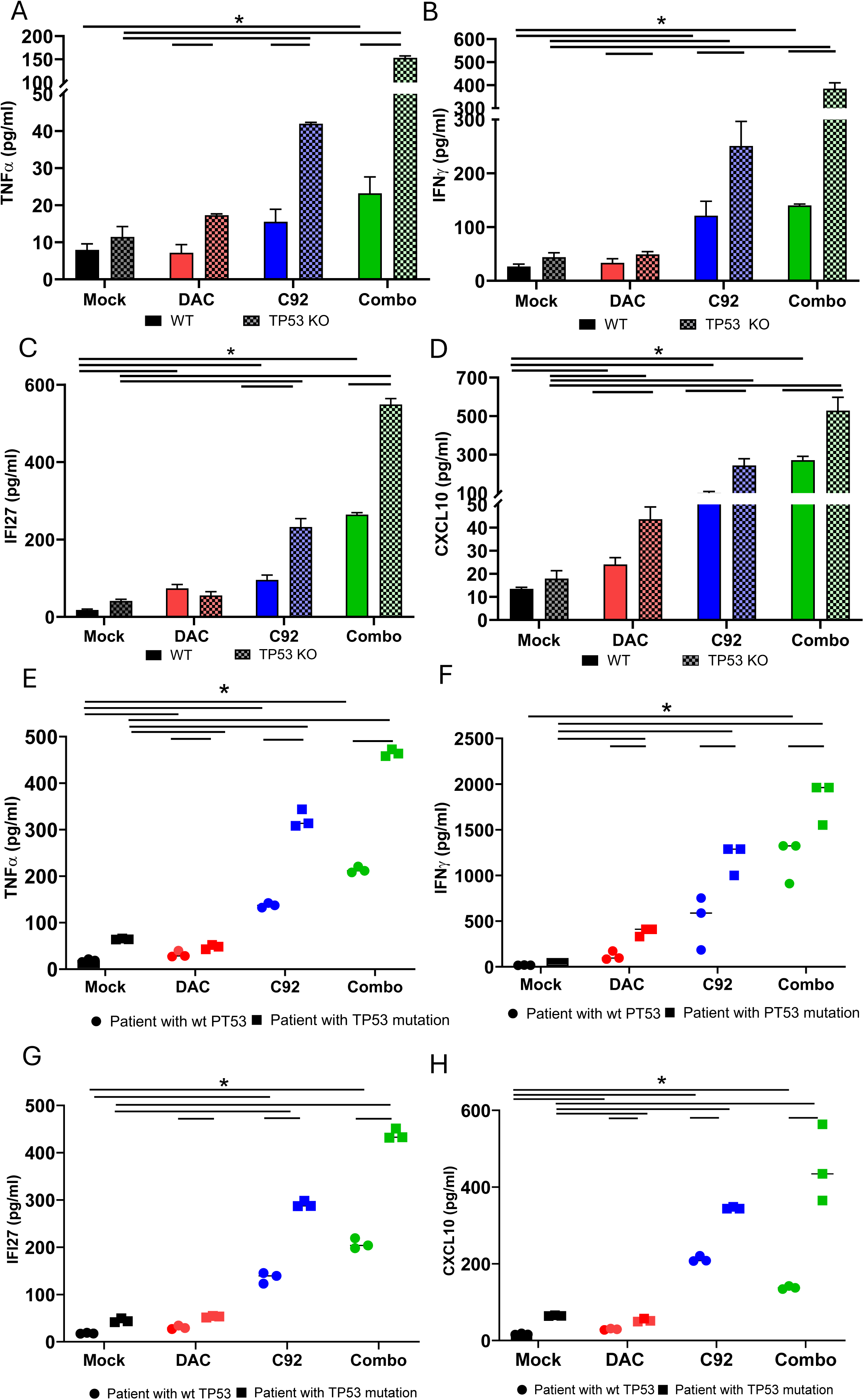
STING agonists C92 in combination with DAC increases inflammasome signaling in primary AML cells and cell lines with mutations or inactivation of TP53. **A-D**. Cytokine expression of **A.**TNF alpha, **B.**IFN gamma, **C.** IFI27 and **D.** CXCL10 respectively assessed by ELISA assay in MOLM-14 WT and MOLM-14 PT53 -/- cells 3 days after DAC (25nM), C92 (250nM), and combination treatment. **E-H.** Dot plot showing cytokine expression of **E.**TNF alpha, **F.** IFN gamma, **G.** IFI27 and **H.** CXCL10 respectively assessed by ELISA assay in primary AML cells WT versus TP53 KO (N=5). All data are presented as mean +/- SEM with p-values derived from two-tailed unpaired Student’s t test or ANOVA as appropriate. * p<0.05, p<0.01, p<0.001, p<0.0001. All experiments were performed at least 3 times.

To begin to understand the genes and molecular pathways affected by the drug treatments, we next tested the transcriptome-wide effects of C92 (250nM) or DAC (25nM) treatment alone or the two drugs in combination, by performing ribosomal-depleted RNA-seq on WT and *TP53* CRISPR KO MOLM-14 cells after 3 days of treatment. As our study queries the impact of drug treatment on genomic perturbation, we first applied gene set co-regulation analysis (GESECA), a pathway analysis method designed to assess multi-conditional data. Both C92- and C92+DAC-treated *TP53* KO cells manifest a conserved positive enrichment of gene sets associated with Type I/II interferon, TNFα, IFNα, IFNγ, and JAK-STAT, with concurrent downregulation of MYC and OXPHOS gene sets (**Fig. 3A-C**). Such down regulation of MYC and its target genes specifically reverses its repression of immune-related signaling in cancer cells (35). Assessment of differentially expressed genes (DEGs) defines a marked increase in interferon-associated cytokines and effector molecules when comparing WT and *TP53* CRISPR KO MOLM-14 cells treated with C92 and C92+DAC. Additionally, in agreement with pathway-level data, MYC is significantly downregulated in *TP53* KO MOLM-14 treated with C92 and C92+DAC (**Fig 3A**). Critically, while the number of DEGs is comparable between C92 and C92+DAC in *TP53* WT cells, there is a clear separation in DEG abundance between these conditions in *TP53* KO cells, with 2429 and 3635 differentially expressed genes in C92 and C92+DAC conditions, respectively (**Fig. 3D**). These observed dynamics confirm that while interferon-associated pathways appear similar by pathway-level metrics, transcriptional induction between C92 and C92+DAC in *TP53* KO cells, in the latter, 39 interferon-associated genes are upregulated more than 1.5-fold between C92 and C92+DAC treatments **(Figs 3B-D)**. Importantly, these combination-specific augmented genes reside in critical interferon gene categories, such as Th1 type chemokines (*CXCL10*, *CXCL11*), RNA detection (*DHX58, EIFAK2, IFIH1, OASL, PARP9, RIGI*), and interferon cell death-associated (*IFI27, IFI44L, IFIT2, MX1, OAS1, OAS2, OAS3, TNFSF10)* genes, as validated by qPCR **(Fig 3E-H)**. Moreover, these analyses reveal orthogonal increases in both overall counts and numbers of differentially expressed repetitive elements (REs), largely restricted to *TP53* KO cells treated with combined C92 and DAC (**SFig 5A, B),** and confirmed by qPCR of a panel of RE elements, including L1PA2, ERV-E4, ERV-L, ERV-K3, ERV-H1 **(Fig 3I-M**, **Table 2**.). Augmented expression of REs in the genome is a key component of DAC induced immune signaling gene expression(12,16).

**Fig 5.**
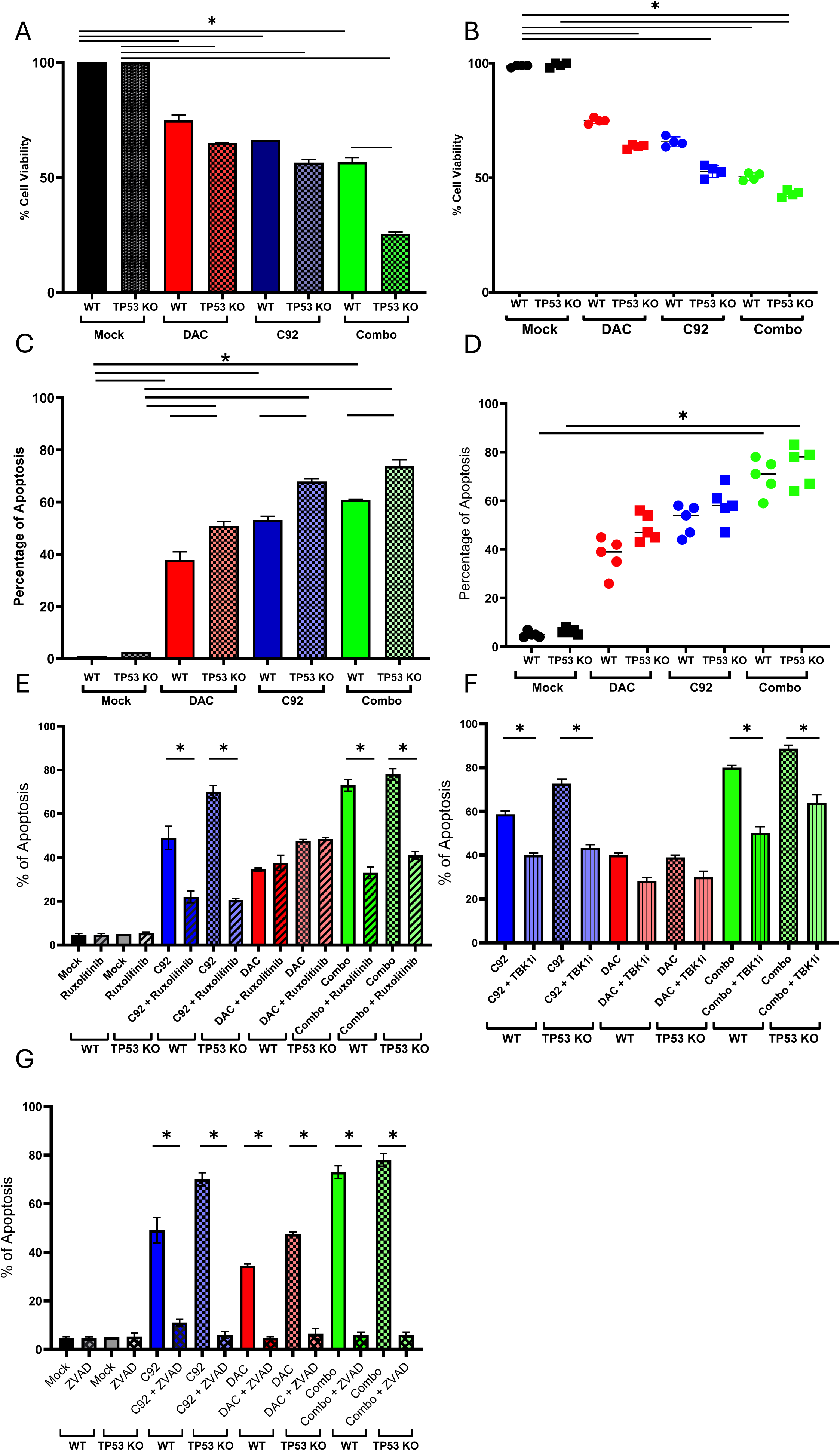
Combination treatment increases apoptosis in *TP53* KO MOLM-14 cells **A.** Bar graph showing cell viability performed by MTS in MOLM-14 WT and MOLM TP53 KO after 3-day treatment with DAC (25nM), C92 (250nM) and combination. **B.** Dot plot showing cell viability performed by MTS in primary AML cells WT versus TP53 -/-. **C**. Annexin V analysis performed via flow cytometry in MOLM-14 WT and MOLM TP53 KO before and after 3-day treatment with DAC (25nM), C92 (250nM) and combination. **D.** Annexin V analysis performed via flow cytometry in primary AML cells. **E.** Annexin V analysis performed via flow cytometry in MOLM-14 WT and MOLM-14 TP53 KO treated to ruxalitinib (JAK inhibitor) in addition to DAC (25nM) and C92 (250nM) and combination therapy. **F.** Annexin V analysis performed via flow cytometry in MOLM-14 WT and MOLM-14 TP53 KO treated with TBK inhibitor in addition to DAC (25nM) and C92 (250nM) and combination therapy. **G**. Annexin V analysis performed via flow cytometry in MOLM-14 WT and MOLM TP53 KO treated with pan-apoptotic inhibitor ZVAD in addition to DAC (25nM) and C92 (250nM) and combination therapy. All data are presented as mean +/- SEM with p-values derived from two-tailed unpaired Student’s t test or ANOVA as appropriate. * p<0.05, p<0.01, p<0.001, p<0.0001. All experiments were performed at least 3 times.

**Table 2.**
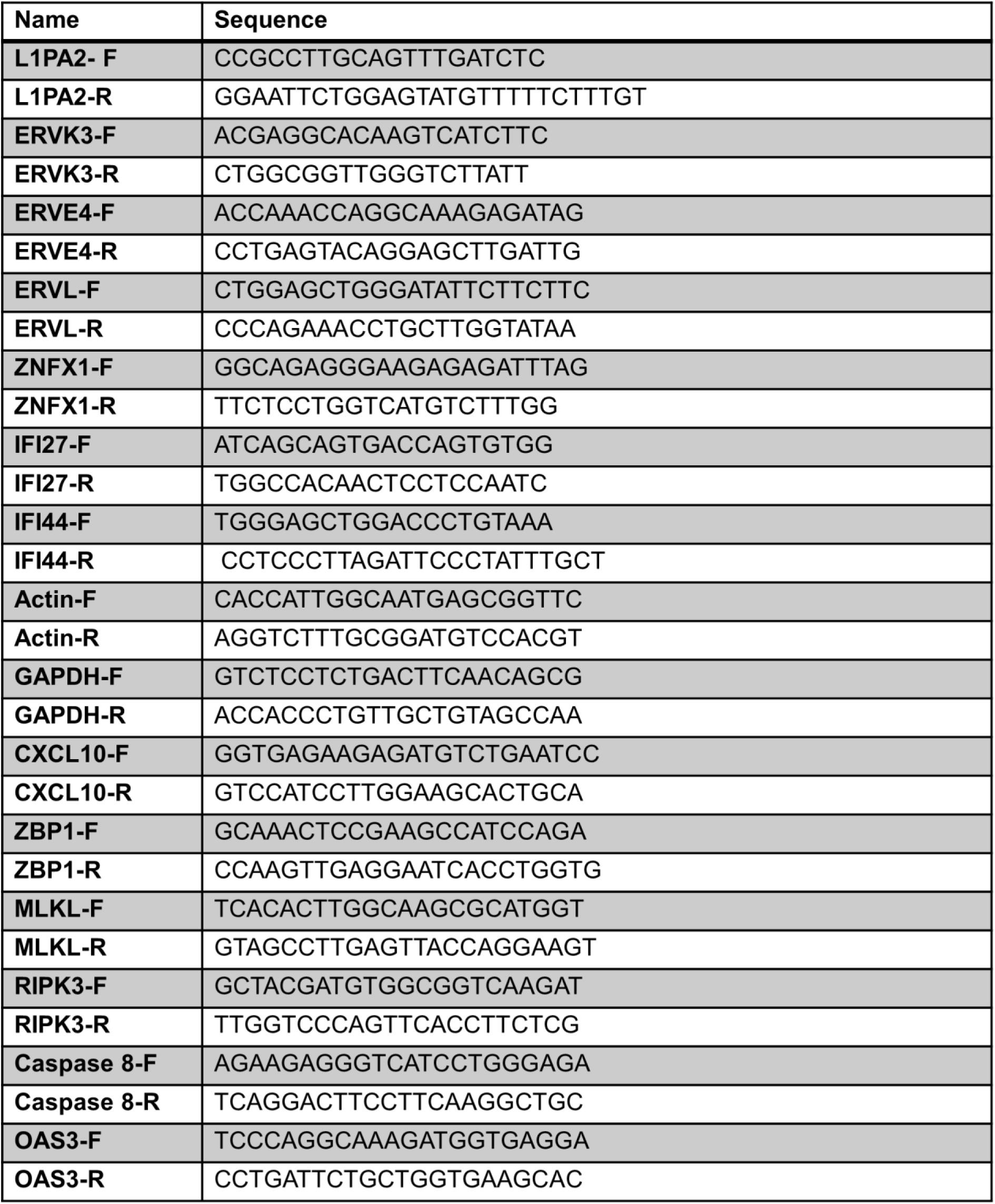
List of primer used to detect gene expression in AML.

Based on the results showing that C92+DAC treatment induces transcriptional increases in individual cytokine and interferon genes associated with Type I/II interferon, TNFα and immune signaling, we assessed by ELISA, CXCL10, TNFα, IFNγ and IFI27 in AML cell lines and primary cells. In MOLM-14 AML cells with WT *TP53*, DAC or C92 treatment alone result in little or no increases in cytokine expression, whereas the drug combination produces significant increases in TNFα, IFNγ, IFI27 and CXCL10 (**Fig 4A-D**). Notably, in *TP53* KO MOLM-14 cells, compared with MOLM-14 parental cells, cytokines are increased by single agent treatment but further increased by the drug combination (**Fig 4A-D**). Similar results are observed in studies of primary AML cells with mutated *TP53* versus WT (N=5) (**Fig 4E-H**, **Table 1**).

Acquisition of this high interferon state in C92+DAC-treated *TP53* KO cells is of particular interest when considering the above concurrent RE induction (**Fig 3**), as this convergence brings substrate, sensors, and effectors together to potentially drive the suicidal interferon-driven cell death response previously noted. Importantly, as assessed with MTS assays, both single agents and the two-drug combination significantly decrease cell viability in both cell lines and primary MOLM-14 WT and *TP53* KO cells (**Fig 5A, B, Table 1**).

To investigate cell death by apoptosis with the above drug treatments, we performed flow cytometric analysis of Annexin V labeling in MOLM-14 and MOLM-14 *TP53* KO cells pre and post drug treatments. C92 or DAC treatment significantly increases apoptosis in MOLM-14 cells, and the combination increases it even further (**Fig 5C**). These results are validated in primary AML cells **(Fig 5D)**. To test whether the apoptotic effects are induced by interferon, we treated these cells with C92 and DAC in the presence of the JAK/STAT interferon inhibitor ruxolitinib. Interestingly, ruxolitinib is reflecting a stronger interferon response to C92 compared to DAC. Moreover, ruxolitinib treatment does not differentially affect *TP53* KO vs WT cells (**Fig 5E**). To further investigate whether STING pathway activation drives apoptosis in AML cells, we examined apoptosis induction by the drug combination in the presence or absence of an inhibitor of the immediate downstream STING target TBK1 (**Fig 5F**). TBK1i treatment abrogates induction of apoptosis by C92 treatment in AML cells, whereas apoptosis induction by DAC treatment is inhibited to a lesser extent. Moreover, TBK1i treatment does not differentially affect *TP53* KO vs WT cells. Lastly, treatment with the pan-apoptosis inhibitor ZVAD rescues all MOLM 14 cells tested from apoptosis induction by C92 and DAC treatment (**Fig. 5G**).

Given that the above apoptotic pathway induced by C92 and DAC treatment was not specifically potentiated in *TP53* KO cells, we questioned whether *TP53*-mutated AML might trigger additional cell death mechanisms. Z-DNA binding protein 1 (ZBP1) is a key innate immune sensor that triggers PANoptosis—an inflammatory cell death pathway combining apoptosis, necroptosis, and pyroptosis. We recently identified a little-studied innate immune gene, ZNFX1, as a master regulator of STING activation in ovarian cancers(36). Cells with high ZNFX1 levels also exhibited concomitant upregulation of the PANoptosis gene ZBP1 (36). Indeed, in our RNA seq data from MOLM-14 *TP53* KO and WT cells treated with C92 and DAC, both ZNFX1 and ZBP1 expression are potentiated in *TP53* KO cells (**Fig 6A**), and validated by qPCR and western blot analyses (**Fig 6B,C**). Moreover, double KO of *TP53* and ZNFX1 in MOLM-14 abrogates this increased expression (**Fig 6D,E**). Since, the PANoptotic pathway encompasses multiple cell death mechanisms, including necroptosis, pyroptosis or apoptosis characterized by increased MLKL, gasdermin D (GSDMD), and caspase 3 expression respectively, we performed qPCR and immunoblotting against these major death pathway signaling proteins, examining for changes in expression and cleavage in *TP53* KO/ *TP53 and ZNFX1* KO versus WT MOLM-14 cells treated with C92 and DAC drug combination. No expression differences are observed in GSDMD, active in the pyroptosis pathway (**SFig 6A,B**). In contrast, qPCR of mRNA showed increases in MLKL transcripts that was abrogated in ZNFX1/TP53 double KO cells (**Fig 6F**). Likewise, a striking increase in cleavage and activation of MLKL steady state protein levels with C92 and DAC treatment was observed in WT and TP53 MOLM14, while cleavage was abrogated in the double KO cells (**Fig 6G-H**). In addition, cleavage induction in phosphorylated Receptor-Interacting-Serine/Threonine-Protein Kinase 3, pRIPK3, upstream of MLKL was observed in WT and TP53 MOLM14 treated with C92 and DAC, while cleavage reduction was detected in double KO cells. Both of these proteins play a crucial role in tumor necrosis factor (TNF)-induced necroptosis (37). Thus, our data suggests that induction of the necroptosis pathway by C92 and DAC treatment is likely regulated by ZNFX1.

**Fig 6.**
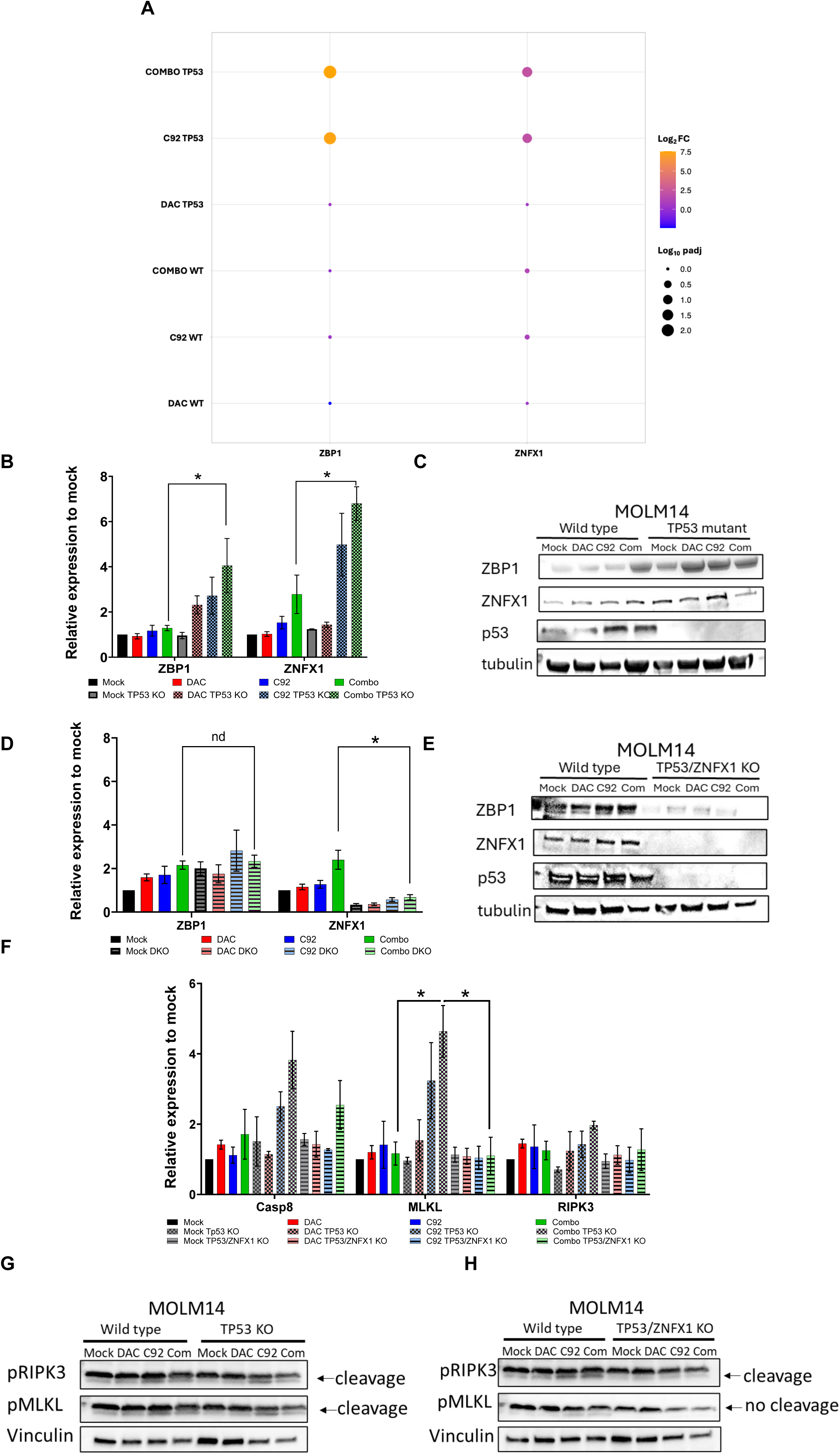
Combination treatment in *TP53* KO MOLM-14 cells activates necroptosis cell death **A.** RNA seq analysis showing ZNFX1 and ZBP1 expression after 3-day treatment with DAC (25nM), C92 (250nM) and combination in MOLM-14 WT and MOLM *TP53* KO. **B.** Validation of ZNFX1 and ZBP1 gene expressions after 3-day treatment with DAC (10nM), C92 (100nM) and combination in MOLM-14 WT and MOLM-14 *TP53* -/- performed by qPCR. **C.** Relative transcript levels of ZNFX1 and ZBP1 in MOLM-14 WT and MOLM-14 *TP53* KO + *ZNFX1* KO (double KO) performed by qPCR following 3 days of treatment with DAC (10nM) and C92 (100nM) and combination. **D.** Western blot depicting protein expression in MOLM-14 WT and MOLM-14 *TP53* -/- after 3 days of treatment with DAC (10nM) and C92 (100nM) and combination. **E.** Western blot depicting protein expression in MOLM-14 WT and MOLM-14 *TP53* KO + *ZNFX1* KO following 3 days of treatment with DAC (10nM), C92 (100nM) and combination. **F.** Relative transcript levels of caspase 8, (40) and RIPK3 performed by qPCR. **G.** Western blot depicting protein expression in MOLM-14 WT and MOLM-14 *TP53* -/- after 3 days of treatment with DAC (10nM) and C92 (100nM) and combination. **H.** Western blot depicting protein expression in MOLM-14 WT and MOLM-14 *TP53* KO + *ZNFX1* KO following 3 days of treatment with DAC (10nM), C92 (100nM) and combination. Protein cleavage and activation depicted by arrows. All data are presented as mean +/- SEM with p-values derived from two-tailed unpaired Student’s t test or ANOVA as appropriate. * p<0.05, p<0.01, p<0.001, p<0.0001. All experiments were performed at least 3 times.

Finally, we queried how all of our above data extrapolates to the *in-vivo* efficacy of our STING agonist C92 and DAC combinatorial approach. We especially examined both human tumor-intrinsic and human host immune responses that might correlate with anti-leukemia drug responses in this setting by utilizing immune-competent AML mouse models. To ask these questions, humanized mice were generated to rule out the previously introduced off target effects of engaging mouse STING signaling. Two- to three-day old NSG-SGM3 immunodeficient mice were humanized by intrahepatic injection of 10^5^ CD34+ stem cells and, at 12 weeks, and data acquired in those screened by flow cytometry for expression of human CD45 and having at least 25% CD45+ cells, a common marker for all human leukocytes **(Fig. 7A,B).** After injection of 0.5 million MOLM-14 cells, and 15 days of treatment with our combinatorial C92 approach, there is significantly more reduction in leukemia burden with the combinatorial drugs than with the single agents alone (**Fig. 7C, D).** Importantly, mouse body weights are not affected by any of the treatments (**Fig. 7E**). Moreover, combination drug treatment significantly increases levels of TNFα, CXCL10, and IFI27 human cytokines in Day 15 plasma samples (**Fig 7F-H**), and IHC analyses of spleen samples for CXCL10 (**Fig. 7I**) and CD8 T cell infiltration of the leukemia microenvironment (LME) (**Fig 7J**). *Our data then suggests that, indeed, C92 in combination with DAC leads to potent anti-leukemic immune responses*.

**Fig 7.**
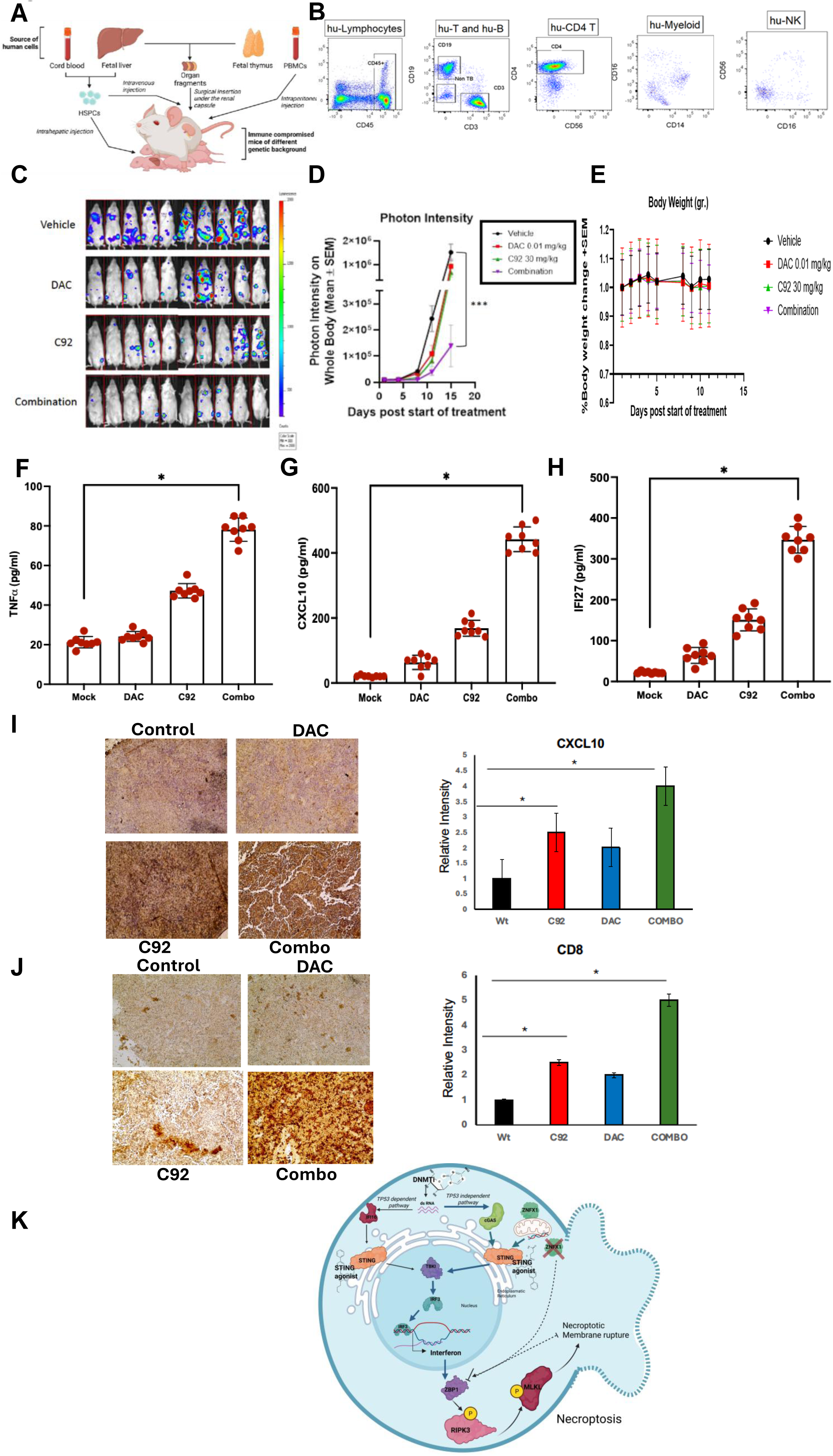
Combination drug treatment decreases tumor growth and increases plasma cytokine levels in humanized mice. **A.** Schematic representation of generation of humanized mice with a functioning immune system. **B.** Flow cytometry showing CD45, CD3, CD56, CD14, and CD16 expression in mice peripheral blood. **C.** Mice imaging showing tumor burden fifteen days after post treatment with DAC (0.01 mg/kg), C92 (30 mg/kg) and combination. **D.** Photon intensity on the whole mouse body fifteen days after post treatment with DAC (0.01 mg/kg), C92 (30 mg/kg) and combination. **E.** Mice body weight fifteen days after post treatment with DAC (0.01 mg/kg), C92 (30 mg/kg) and combination. **F-H**. Cytokine expression of **F.** TNFα, **G.** IFNγ, and **H.** CXCL10 respectively assessed by ELISA assay in plasma from retroorbital bleeds of euthanized mice 15 days after DAC (0.01 mg/kg), C92 (30 mg/kg), and combination treatment. **I-J.** Immunohistochemistry performed on spleen tissue showing **I.** CXCL10 cytokines and **J.** CD8 T cell infiltration respectively 15 days after DAC, C92, and combination treatment. **K.** Graphical model showing Necroptosis pathway activation by DAC and C92 STING agonist combination therapy in AML cells depleted of TP53. DAC and C92 combination therapy demonstrate increased TP53 independent STING pathway that leads to interferon transcription and triggers a ZNFX1-ZBP1 PANoptosis cell death pathway. Activated ZBP1 along with RIPK3 and MLKL phosphorylation are key features of necroptosis, with membrane rupture occurring as a part of this cell death pathway. In cell lines where ZNFX1 is depleted, ZBP1 is completely abrogated and necroptosis does not occur. All data are presented as mean +/- SEM with p-values derived from two-tailed unpaired Student’s t test or ANOVA as appropriate. * p<0.05, p<0.01, p<0.001, p<0.0001. All experiments were performed at least 3 times.

## Discussion

*TP53-*mutated AML remains a clinical challenge and an unmet need (7). Here, we show that a therapeutic barrier imposed by these mutations can be overcome by direct activation of the cGAS/STING pathway, using a novel human-specific STING agonist, C92. This new drug binds all variants of human STING specifically and causes allosteric changes that stabilize the activated dimer (21). Notably, we show for the first time that combining C92 with epigenetic drugs produces synergistic cytotoxicity through activating the STING-dependent interferon-inflammasome pathway in *TP53*-mutated AML cell lines and primary cells. Importantly, our data show that this drug combination drives ZBP1/ZNFX1-dependent necroptotic cell death in *a TP53-independent* manner and represents a potentially promising novel therapy for *TP53*-mutated AML (**Fig 7K, model**). Apoptotic cell death in *TP53*-mutated AML has previously been linked to STING activation. Combining clinically-relevant STING agonists with BH3-mimetic drugs was reported to efficiently kill *TRP53*/*TP53*-mutant mouse B lymphoma, human NK/T lymphoma, and AML cells(19). Our data shows that while the DNMTi-STING agonist combination induces apoptosis, it is not specifically potentiated in *TP53-*mutated vs WT AML cells. In contrast, we show for the first time that the innate immune gene ZNFX1, which we recently reported as a master regulator of mitochondrial dysfunction and STING activation(36), is implicated in driving ZBP1, a key player in PANoptosis, in AML cells with *TP53* KO versus WT. ZNFX1 appears to be required for expression of the innate immune and PANoptosis gene ZBP1(38), known to regulate three major inflammatory cell death pathways: 1) RIPK3-caspase-8 complex GSDMD-mediated pyroptosis, 2) caspase-3/caspase-7-mediated apoptosis, and 3) MLKL-mediated necroptosis. Our data show that C92 and DAC treatment specifically mediates RIPK3 and MLKL-associated necroptotic cell death in a *TP53*-independent manner. Moreover, since these cell death mechanisms are regulated by ZNFX1, this mechanism is likely to involve mitochondrial dysfunction, as suggested by reduced expression of OXPHOS genes in our RNA seq analysis, and as we have previously reported in ovarian cancer (36). More importantly, necroptosis is an intensely inflammatory form of cell death, releasing danger-associated molecular patterns (DAMPs) that alert the immune system, potentially turning “cold” tumors into “hot” tumors that are more vulnerable to immunotherapies like checkpoint inhibitors (39–41). These studies then predict that combinations of STING agonists with immune checkpoint inhibitors may be efficacious for patients with TP53-mutated AML, previously not responding to immune therapy.

All of the above dynamics are tightly associated with presence of WT versus mutant *TP53* in maintaining stability of the epigenetic program in cells(6), playing a role in silencing repetitive DNA sequences, REs and noncoding RNAs in the genome(42). In the absence of TP53, DNA hypomethylation leads to a marked increase in transcription of these repeats. This phenomenon has been termed TRAIN (Transcription of Repeats Activates INterferon) and suggests a mechanism by which TP53 and the interferon system collaborate to prevent the accumulation of cells with “unleashed” repeats (42). In preclinical settings, *TP53*-mutated AML cells show sensitivity to DNMTis(5,6). Clinically, *TP53*-mutated AML has higher response rates to DNMTis than to chemotherapy, but responses are not durable (8,43). Our studies show for the first time in the setting of *TP53* KO AML, that C92 treatment alone and in combination with DNMTis augments expression of genes residing in critical IFN gene categories, including Th1 type chemokines (*CXCL10*, *CXCL11*), RNA detection (*DHX58, EIFAK2, IFIH1, OASL, PARP9, RIGI*), key IFN induced viral mimicry (*IFI27, IFI44L, IFIT2, MX1, OAS1, OAS2, OAS3, TNFSF10)* and cell death-associated genes. All of our data then supports the intriguing hypothesis that in the absence of TP53, DAC and C92 combination treatment increases transcription of RE’s, leading to marked upregulation of IFN-driven cell death.

How loss of p53 function in AML impacts the kinetics of STING activation by synthetic ligands is an important open question. In the basal state, STING resides on the ER where it binds ligands to trigger a sequence of post translational modifications and enters a highly regulated Coat Protein Complex I (COPI) dependent vesicle trafficking through the Golgi to a post Golgi membrane (44,45). At this latter site, STING is phosphorylated by TBK1 to function as a scaffold for the activation of IRF3. Multiple mechanistic hypotheses explaining TP53 regulation of STING are possible. A direct physical interaction between TP53 and STING is unlikely as multiple proteomic interactome studies using STING have failed to identify TP53 as a STING binding partner (46,47). For example, STING activation is highly sensitive to cholesterol content of the ER membrane (48) and TP53 regulates the Sterol O-Acyltransferase-1 (SOAT) enzyme involved in the esterification and storage of ER cholesterol (49,50). STING activation is governed by the STIM1 ER Ca^2+^ sensor (51), and WT TP53 directly binds to the sarco/ER Ca^2+^-ATPase (SERCA) pump at the ER, changing its oxidative state, leading to an increased Ca^2+^ load (52). Thus, studies of the ER in *TP53*-mutated and -WT cells are required to elucidate these mechanisms.

In summary, our present data support the potential of next-generation STING agonists for providing therapeutic efficacy against AML harboring *TP53* mutations. This efficacy may well be translated to multiple other *TP53-*mutated cancers, such as ovarian, endometrial and lung. In addition to evaluation of C92 as a single agent, our data support initiation of trials for combining STING agonist with epigenetic drugs in patients with *TP53*-mutated AML. We are entering an exciting era of studies which will undoubtedly come to include additional strategies for utilizing next-generation STING agonists such as C92, and particularly for enhancing efficacy of other immune therapies. This latter approach should be tested in treatment-resistant AML and other cancers.

## Acknowledgements

We thank Curadev Pharma and in particular CEO Arjun Surya and CSO Monali Banerjee for generous donation of STING agonist drug C92 and for important insightful discussions. Our studies were supported by the National Cancer Institute–Cancer Center Support Grant P30 CA134274 University of Maryland Marlene and Stewart Greenebaum Comprehensive Cancer Center (F.V.R., R.L, B.C, M.R.B); the Human Genetics Graduate Program, University of Maryland (Z.G.); the Maryland Department of Health’s Cigarette Restitution Fund Program (F.V.R., M.R.B.); Specialized Program of Research Excellence (SPORE) program, through National Cancer Institute (NCI) grant P50CA254897 (L.S., K.T., M.A., F.V.R, S.B.B, M.J. T., K.P.N, Z.G. A.T., A.H.); and the Van Andel Research Institute – Stand Up To Cancer Epigenetics Dream Team (S.B.B., M.B.R., F.V.R., K.P.N.). The indicated Stand Up To Cancer Grant is administered by AACR, the scientific partner of Stand Up To Cancer. M.J.T. is a recipient of The Evelyn Grollman Glick Scholar Award. The content is solely the responsibility of the authors and does not necessarily represent the official views of the NIH.

## Authorship Contributions

K.T., L.S., S.B.B., D.P., A.H., M.J.T, M.R.B., and F.V.R. designed research; K.T. L.S, Z.G, M.A., B.C., R.G.L., A.T., G.S. performed research and analyzed data; K.T., L.S., D.P., A.H., M.J.T., S.B.B., K.P.N, M.R.B. and F.V.R. wrote the paper.

