## supplemental fig for "STING agonists in combination with epigenetic drugs potentiate ZNFX1-driven inflammatory necroptosis in *TP53*-mutated AML"

### SFig 1

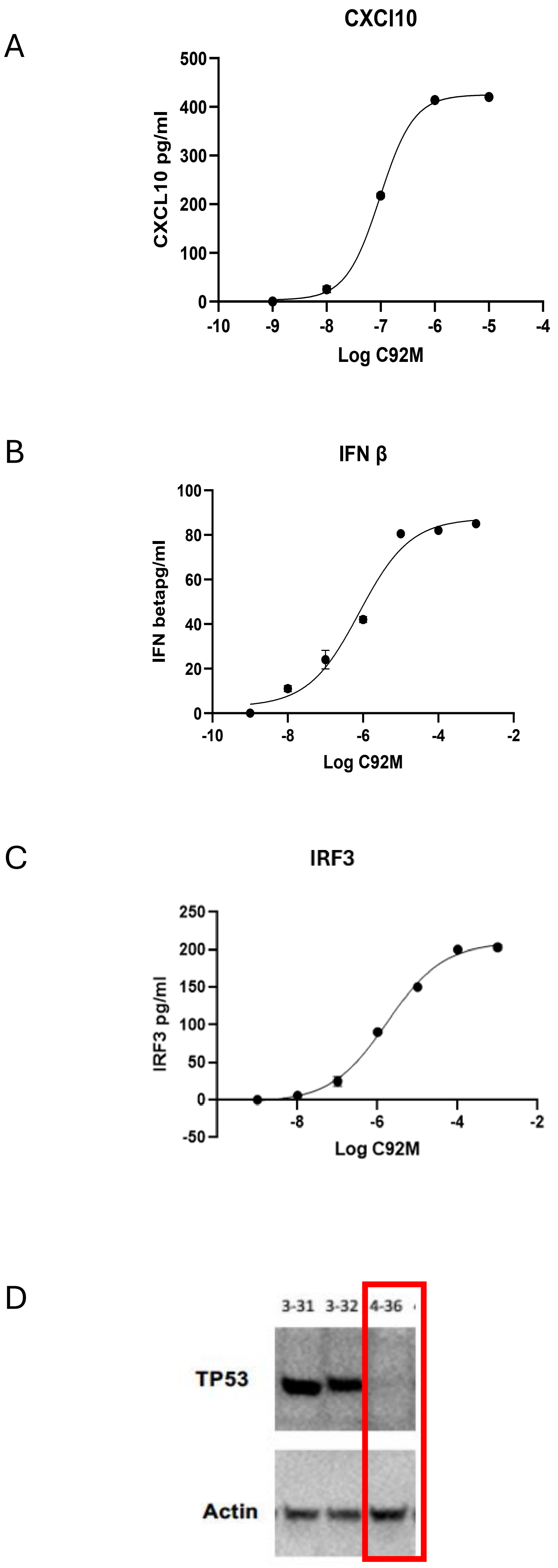


**SFig 1.** STING agonist C92 activates STING in a TP53-independent manner. **A-C**. Expression of proinflammatory cytokines CXCL10, IFN beta, and IRF3, respectively in MOLM-14 measured by ELISA. **D.** Western blot of TP53 in isolated clones, 3-31, 3-32 and 4-36 from CRISPR CAS KO of TP53 in MOLM14 AML. Red box identifies clone 4.34 as TP53 KO. Actin was used as a loading control.

SFig 2

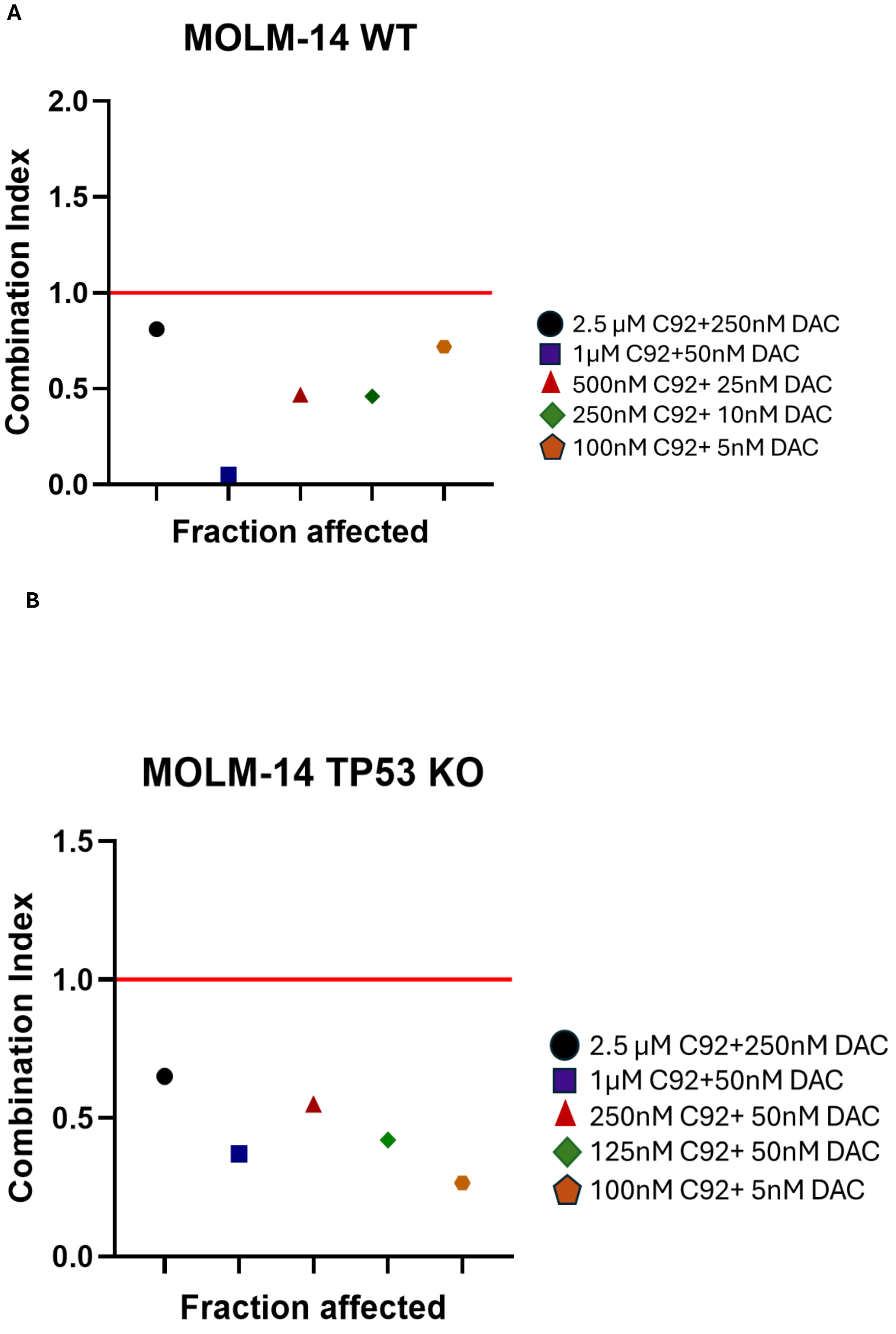


**SFig 2. C92 and DAC are synergistic in AML cell lines MOLM14 and MOLM14 *TP53* KO.** The synergistic relationship between C92 and DAC in cell viability assay (MTS) after three days of treatment. CompuSyn analysis plotted with GraphPrism of combination index (CI) in fraction of cells affected (Fa) is shown in A. MOLM14 wild type and **B.** MOLM14 *TP53* KO AML cells.

**SFig 3**

**A**

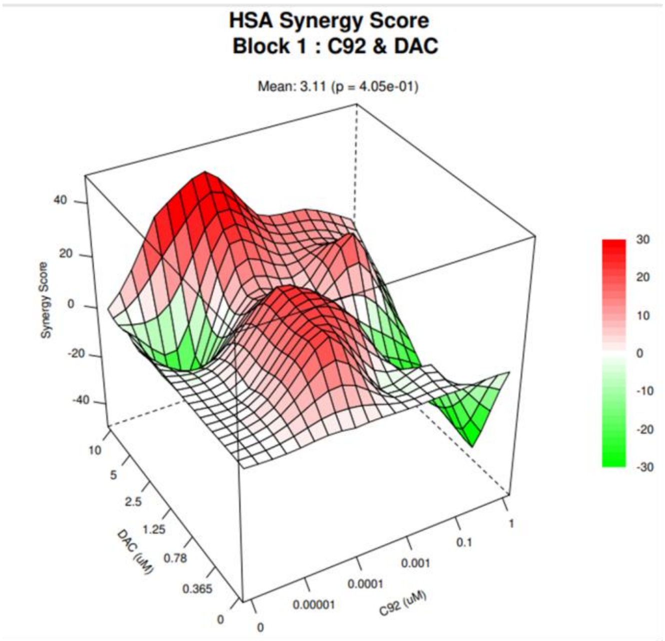


**B**

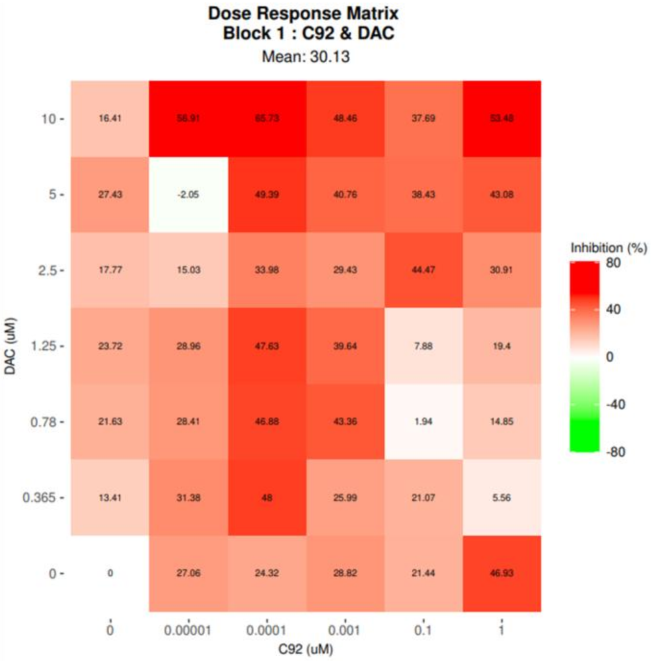


**C**

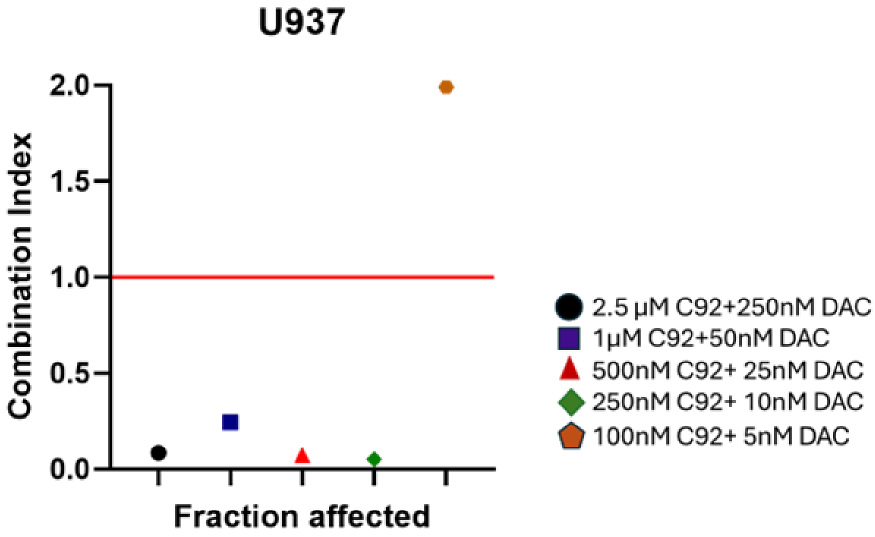


**SFig 3. C92 and DAC are synergistic in AML cell line with TP53-mutated AML. A-B.** The synergistic relationship between C92 and DAC in cell viability assay (MTS) after three days of treatment in AML cell line U937 with *TP53*-mutated. **A**. The Synergy Finder Plus tool was used to visualize drug interaction effects following exposure to a concentration matrix of decitabine (DAC) and C92 in U937**. B.** Heatmap visualization and synergy scores were generated using HSA model. **C.** CompuSyn report and plot of combination index (CI) in fraction of cells affected (Fa) is shown in U937

**SFig 4.**

A.

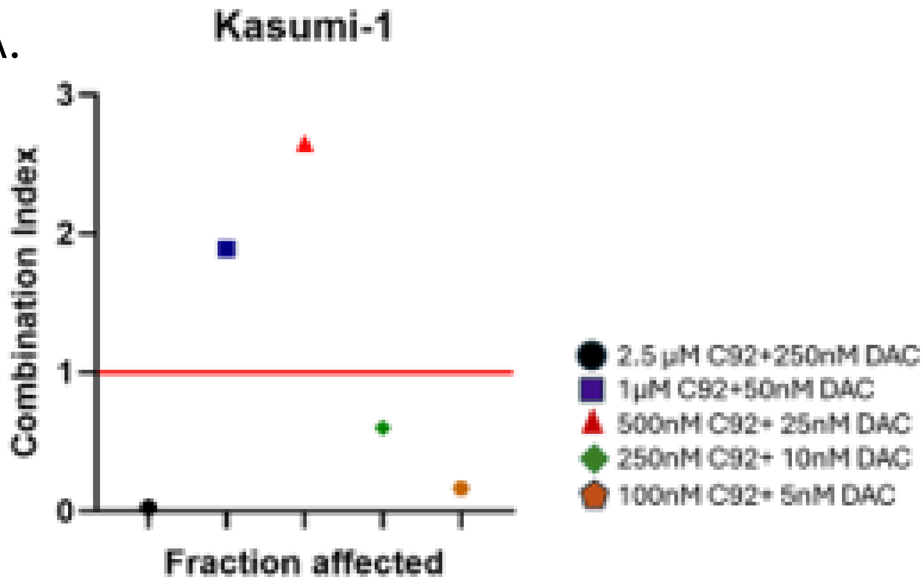


B.

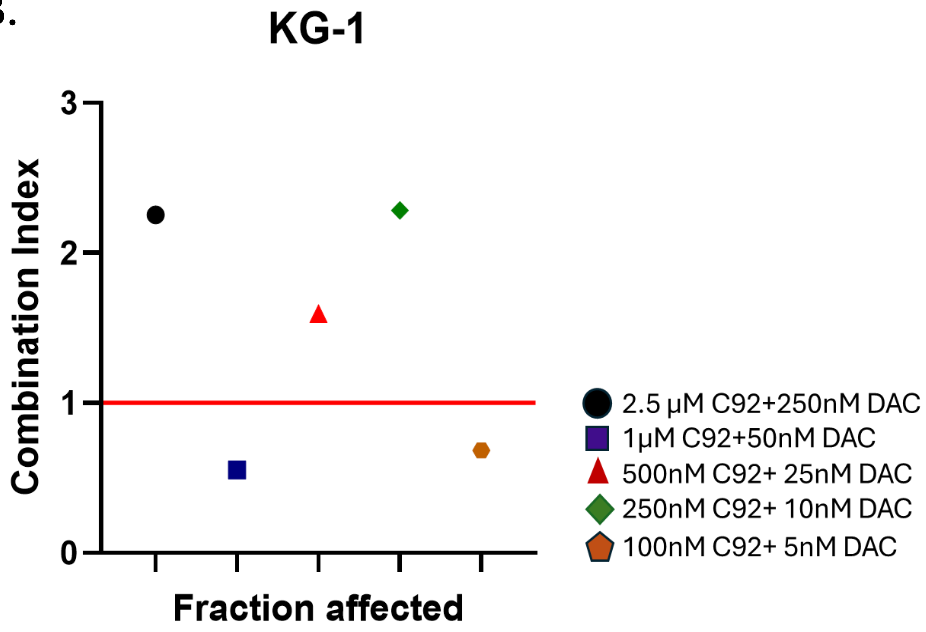


**SFig 4. C92 and DAC are synergistic in AML cell line with TP53-mutated AML. A-B.** The synergistic relationship between C92 and DAC in cell viability assay (MTS) after three days of treatment in AML cell line **A.** Kasumi-1 and **B.** KG-1 **A**.

**SFig 5**

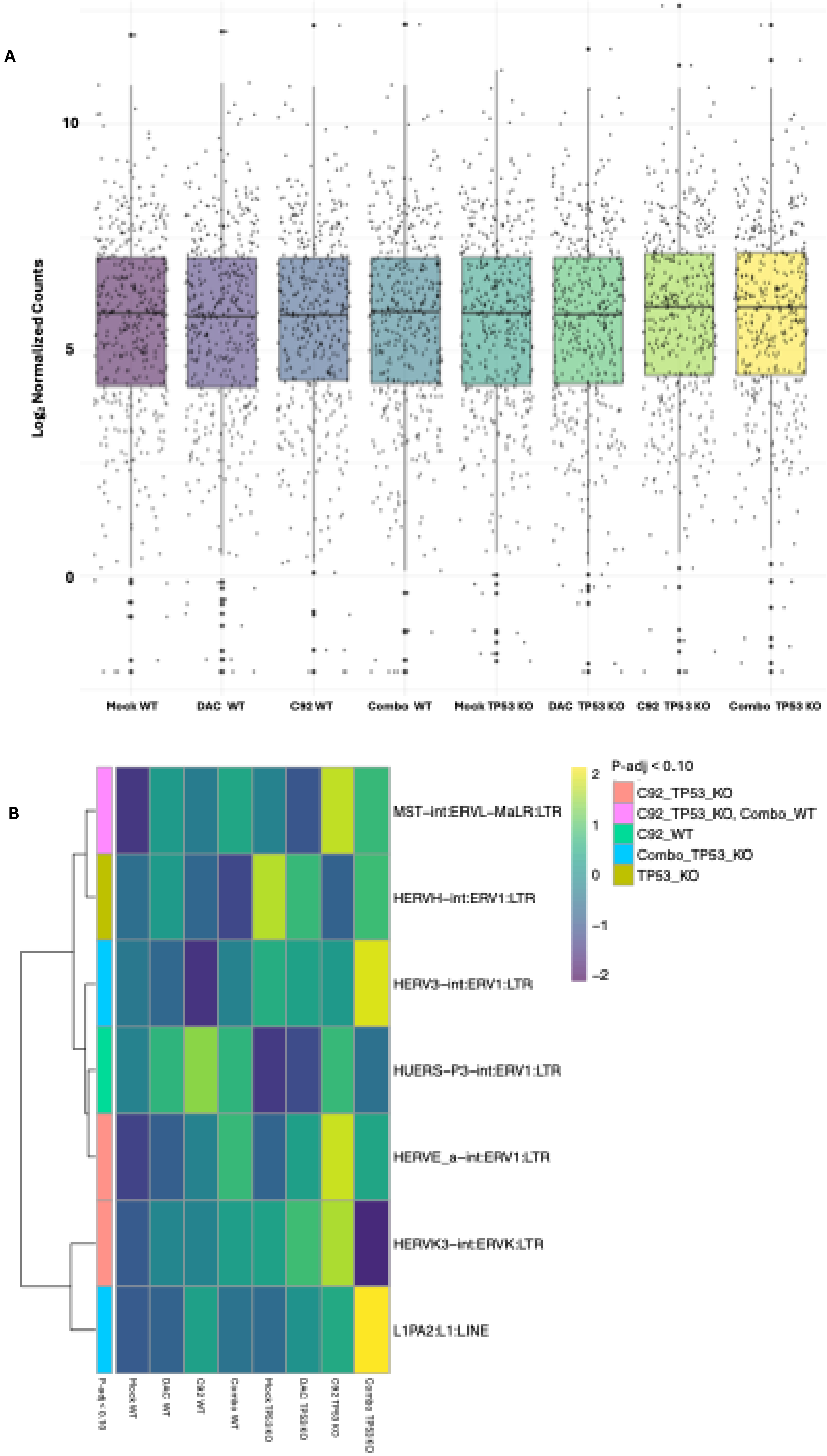


**SFig. 5: *TP53* KO potentiates repetitive element induction from STING agonist (CRD3874). +/- Dac**
**(A)** Boxplots of normalized counts derived from Tetranscripts->DESeq2 pipeline applied to n=2 averaged RNA-seq datasets collected following 3-day treatments (conditions indicated in the plot) in MOLM14 *TP53* wild-type and MOLM14 *TP53* KO cells . Bold line at center represents median for dataset, individual dots are annotated GENCODE repetitive elements. (**B**) Scaled heatmap of median normalized counts for selected repetitive elements based on statistical significance determination by DESeq2 in at least 1 dataset. Dataset with statistically significant result indicated in the plot by padj<0.10 annotation. Hierarchical clustering of genes by Ward.D2**.**

### SFig 6

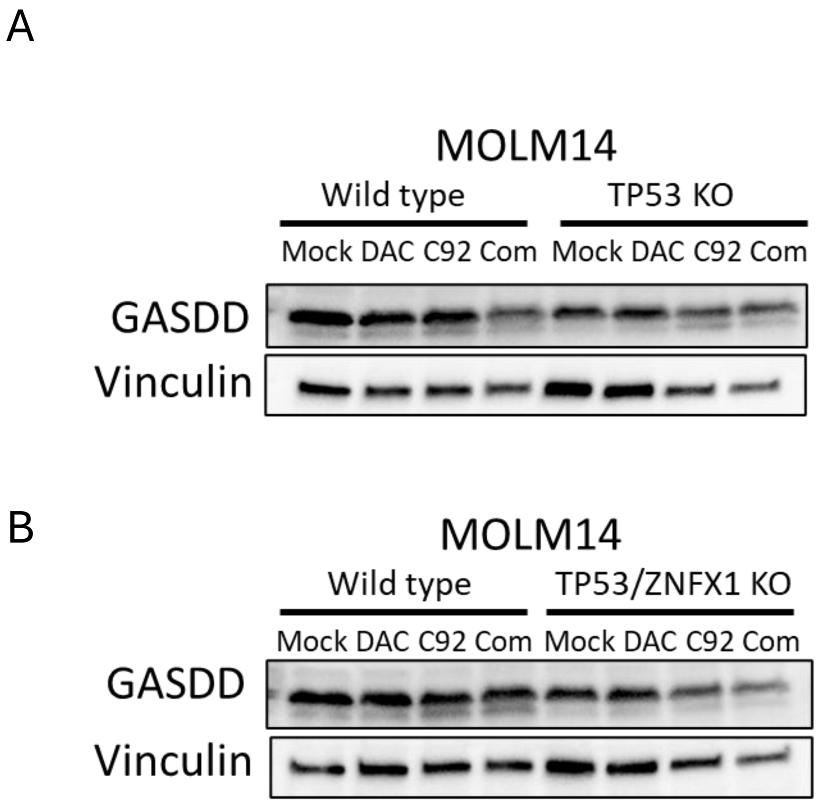


**SFig 6.** Western blot depicting protein expression of gasdermin D in **A.** MOLM-14 WT and MOLM-14 TP53 KO and **B.** MOLM-14 WT and MOLM-14 TP53 KO + ZNFX1 KO following 3 days of treatment with DAC (10nM), C92 (100nM) and combination. Vinculin was used as loading control.
